# Spatio-Genetic and Phenotypic Modelling Elucidates Resistance and Re-Sensitisation to Treatment in Heterogeneous Melanoma

**DOI:** 10.1101/463877

**Authors:** Arran Hodgkinson, Laurent Le Cam, Dumitru Trucu, Ovidiu Radulescu

## Abstract

Although novel targeted therapies have significantly improved the overall survival of patients with advanced melanoma, understanding and combatting drug resistance remains a major clinical challenge. Using partial differential equations, we describe the evolution of a cellular population through time, space, and phenotype dimensions, in the presence of various drug species. We then use this framework to explore models in which resistance is attained by either mutations (irreversible) or plasticity (reversible). Numerical results suggest that punctuated evolutionary assumptions are more consistent with results obtained from murine melanoma models than gradual evolution. Furthermore, in the context of an evolving tumour cell population, sequencing the treatment, for instance applying immunotherapy before BRAF inhibitors, can increase treatment effectiveness. However, drug strategies which showed success within a spatially homogeneous tumour environment were unsuccessful under heterogeneous conditions, suggesting that spatio-environmental heterogeneity may be the greatest challenge to tumour therapies. Plastic metabolic models are additionally capable of reproducing the characteristic resistant tumour volume curves and predicting re-sensitisation to secondary waves of treatment observed in patient derived xenograft (PDX) melanomas treated with MEK and BRAF inhibitors. Nevertheless, secondary relapse due to a pre-adapted subpopulation, remaining after the first wave of treatment, results in a more rapid development of resistance. Our model provides a framework through which tumour resistance can be understood and would suggest that carefully phased treatments may be able to overcome the development of long-term resistance in melanoma.

## 1. Introduction

### 1.1. Mathematical Background

The vast majority of existing, quantitative models of drug resistance are based on discrete stochastic mechanisms of evolution, which fail to take into account the intermediary stages and continuous nature of phenotypic development [1, 2, 3, 4].

Of the continuous models, several provide insights into the dynamics of evolutionary processes but are often restricted to single cell or non-spatial population models [5, 6], necessarily containing space averaging assumptions (well-stirred reactor hypothesis). Of these models, few take into account the prominent theory of PE [7] or have the depth to explain its significance in the context of drug resistance. Herein, we present a continuous spatio-structuro-temporal model to describe both the dynamics of the population of evolving tumour cells as a whole and how targeted therapy can produce resistant strains. We further use the model to recommend future strategies for prevention of this process.

One recent study has further looked at the effect of diffusion-based drug gradients on the effective outcome of population diversity and heterogeneity [8]. This heterogeneity is evident in the biological literature but is yet to be explained by existing mathematical models.

A new addition to the variety of available bio-mathematical modelling frame-works has been spatio-structuro-temporal modelling, introduced by Domschke *et al.* [9] and later subjected to higher-dimensional simulation and numerical analysis [10]. This allows one to represent not only the spatial aspects of a population but also, simultaneously, some underlying aspect of its structure, giving one more insight into the co-evolution of these characteristics. This model has since been extended further [11] but has not yet been used to look at intrinsic properties of tumours, with respect to their systematic resistance to targeted therapies.

### 1.2. Biological Motivation

Current mathematical abstractions of the biological paradigm for drug resistance characterise the biological system as existing in a series of discrete states; perhaps susceptible cells, cells with resistance to drug 1, cells with resistance to drug 2, and cells with resistance to both drugs. This discrete interpretation (Fig. 1a), however, is not born out in experimentation since the observation of cells under the influence of any given drug will demonstrate a spectrum of response patterns. The common assumption that cells instantaneously realign themselves to a ‘resistant’ phenotype also appears to presuppose the eventual survival of such cells. Moreover, gene expression levels of a given cell population submitted to treatment do not appear to exhibit strong qualitative differentiation and are more accurately described as a continuum.

**Figure 1:**
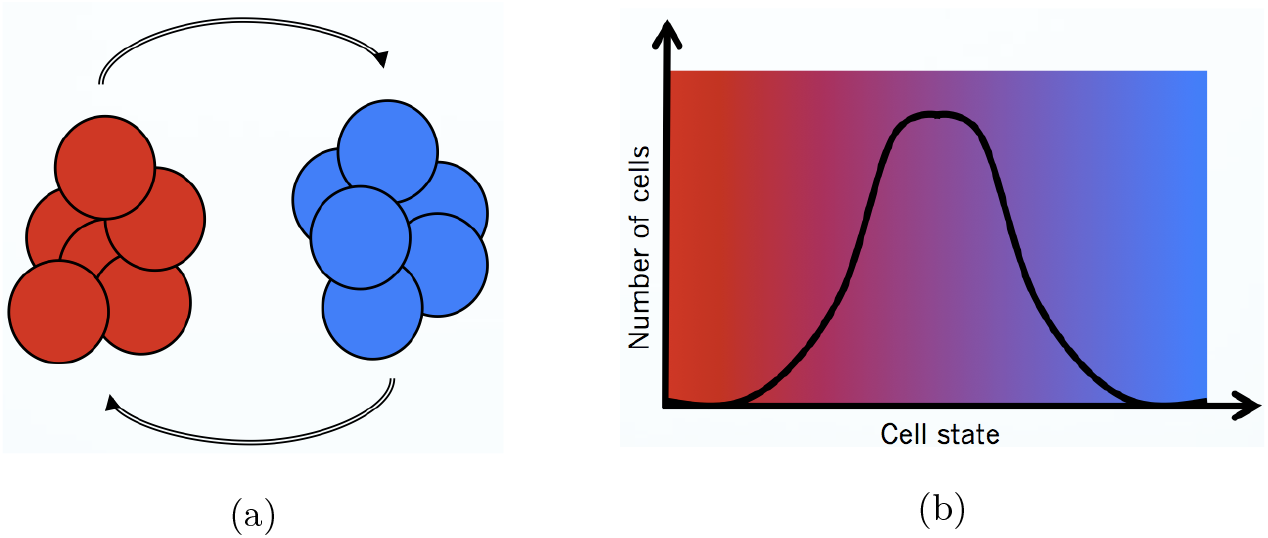
Diagrams indicative of (a) a discrete paradigm, wherein a cell undergoes instantaneous and complete transitions between healthy and resistant states, and (b) a continuous paradigm, in which a cell undergoes a continuous alteration between two extreme healthy and resistant states and is capable of inhabiting all points between these two extremes.

Therefore, we take an alternative approach to modelling wherein we consider the cellular population as a single population which is continuously variable through some structural dimension, as opposed to previous discretised descriptions (Fig. 1b). The structural dimension can be understood as a set of variables characterizing the cell state at a molecular and/or phenotypic level. Within the cell population, subgroups are differentially sensitive to drugs and may exhibit differing proliferative and migratory behaviours, more generally. This gives us extended scope to model the more nuanced aspects of the heterogeneous cellular pathways towards resistance and invasion of the surrounding tissue in cancer.

In the proceeding section (Section 2), we present a general mathematical approach to modelling biological cancer systems whose spatial and structural dynamics are coupled and introduce the various terms within this system. Moving forward we provide a possible application for this model in the study of systems who develop resistance through the sequential mutation of particular oncogenes and the effects of BRAF inhibitors and immunotherapies on this development (Section 3). The results for this mutational model are then studied in detail, with particular interest given to the effects of the order and methodology of treatment and heterogeneity in the tumour environment (Section 4). Next, we provide a second possible application for this model in the study of systems whose metabolism of certain nutrients, particularly the metabolism of glucose through glycolysis or oxidative phosphorylation, shapes their response to drugs, resulting in a plastically resistant system (Section 5). We then explore results coming from this metabolically plastic system with a specific view to understanding the effect of treatment of spatial and metabolic heterogeneity and the resulting responses to treatment (Section 6). Finally we discuss the results from both of these systems in the wider contrast and the ramifications of this current study (Section 7).

## 2. Presentation of the General Model

Herein, we present a mathematical model that contains

(1) One cell species function, denoted *c*(*t*, *x*, *y*), depending on time *t*, space *x*, and structure *y*, representing a continuous distribution of mutational or metabolic phenotypes of cancerous cells:

(1a) The structure variables *y* describe either mutational or metabolic status of the cell. In general, cells will be able to move in either a positive or negative mutational or metabolic direction, depending on the paradigm in question and possibly based on environmental factors.
(1b) The mutational or metabolic alterations taking place within this cellular species will fundamentally alter its behaviour and the nature of its interaction with the micro-and macro-environments,
(2) A function representing the extracellular nutritional environment (ECNE), denoted *υ*(*t*, *x*); including the collagen matrix, distributed fibronectin, and vasculature, assumed to be proportional to one another as explored in the mathematical model of Gatenby [12],
(3) A vector valued function representing the concentrations of diffusible molecular species, denoted *m̄*(*t*, *x*), including metabolites, metallo-proteases, chemo-attractants or chemo-repellents, which will have the ability to mediate the interactions among the variables *c*(*t*, *x*, *y*), and ECNE, *υ*(*t*, *x*),
(4) A vector valued function representing the concentrations of some medicines, denoted *p̄*(*t*, *x*), of detriment to the growth of certain of the cancerous species.

In the following, we describe the main steps for building the model. Details of the mathematical approach to establishing this system of equations may be found in Appendix A.

Mathematically, we employ a multi-dimensional framework which allows for the coupling of spatial dynamics, in *x*, with other biological or biochemical dynamics in the cells themselves, which we call structural dynamics and denote by *y*. Then we can use an existing mathematical framework [9, 11] to deduce that the change in cell density *c*(*t*, *x*, *y*) is given by the continuity equation

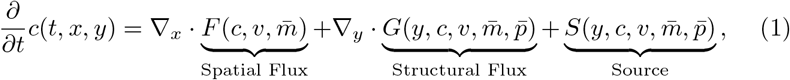

where *ā*·*b̄* stands for the dot product of vectors *ā* and *b̄*.

Through this, we recognise that the function *F* (*c*, *υ*, *m̄*) describes the movement of the cellular population in space, whilst *G*(*y*, *c*, *υ*, *m̄*, *p̄*) describes the structural change in the cellular population, and *S*(*y*, *c*, *υ*, *m̄*, *p̄*) describes the overall change in the population or the number of cells entering or leaving the system through mitosis or apoptosis/necrosis, respectively.

### 2.1. Spatial flux of the cellular population

Begin by denoting *ρ* the collective spatial volume of the cellular and ECNE populations, defined as

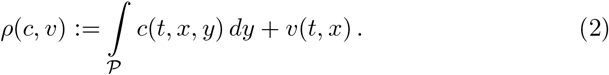

where *𝒫* is the structural domain. This *ρ*(*c*, *υ*) then represents a measure of the total volume occupied by the cellular and ECNE population, together, and will allow us to model the unoccupied volume into which the cells and ECNE may grow. Further, we assume that the cell spatial dynamics are given by diffusion, chemo-and haptotactic directed transport, as in Chaplain *et al.* [13]. Diffusive dynamics correspond to autonomous stochastic motility in spatial cellular dynamics whilst chemo-and haptotaxis correspond to directed motion evoked through attraction to biochemicals or substrate components, respectively. The diffusion, chemotactic, and haptotactic rate constants are then given by *D_c_*, *χ̄_m_*, and *χ_υ_*, respectively. This may be mathematically represented as the following term:

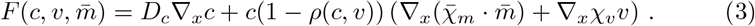

As in [13], the chemo-and haptotactic fluxes are volume constrained and vanish when the collective spatial volume reaches a maximum capacity that, without loss of generality, is considered equal to one. A simple way to take this constraint into account is to consider that these two fluxes are proportional to 1 − *ρ*, where *ρ*(*c*, *υ*) is defined as in (2).

### 2.2. Structural flux of the cellular population

The structural flux is the sum of two terms, an advection flux and a structural diffusion flux, corresponding to biased and unbiased evolution in the structure space, respectively.

In order to define the advection flux we introduce the function Ψ(*y*, *m̄*, *p̄*), representing the normalized structural velocity, who is dependent upon the population’s structural distribution, the local nutrient concentration and the local concentration of drugs. Given some maximal rate for the population’s velocity through the structural dimension, *r_μ_*, the structural velocity shall be given by *r_μ_*Ψ(*y*, *m̄*, *p̄*), where the normalized structural velocity satisfies |Ψ(*y*, *m̄*, *p̄*)| ≤ 1. The structural advection flux term is the product of the structural velocity and the cell distribution density and reads

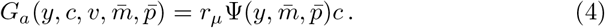

In this paper, we shall consider one cellular system in which behaviour is adapted through the accumulation of consecutive mutation (Section 3) and one in which a cell may plastically evolve its behavioural phenotype through metabolic reprogramming dynamics (Section 5). For each of these scenarios, it will be necessary to define a distinct and biologically relevant form for the function Ψ(*y*, *m̄*, *p̄*).

Diffusion in structural space can occur as the result of a stress, following a change of environmental conditions. In order to adapt to the environment, the population tends to diversify its behaviour which leads to an increase in spread of the *y*-space cell distribution. This diversification of behavior can be phenomenologically described by a structural diffusion matrix Σ(*y*, *m̄*, *p̄*). The structural diffusion flux is supposed to satisfy Fick’s law and reads

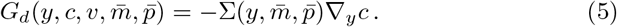

Although structural diffusion is possible both in a mutational and a metabolic context, in this paper we will consider it only in relation to metabolic remodeling.

The total structural flux is the sum of the structural advection and structural diffusion terms and reads

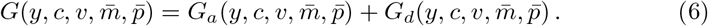

### 2.3. Source/Sink terms for the cellular population

The growth of any given cell will be dependent on an assortment of intracellular and environmental factors, including its structural state, *y*; the availability of nutrients, *m̄*; and the volume surrounding the cell which has not yet been filled, *ρ*(*c*, *υ*). Therefore, we write the growth rate of the population generically as Φ(*y*, *m̄*, *c*, *υ*) such that we may define its particular dynamics for the considered scenario. It is important to remember that this term accounts only for growth of the cell population and not the negative growth caused by the introduction of drugs.

It is clear that, since drugs are typically designed to exploit a particular behaviour or dependence of a given cancerous population, its effectiveness will be dependent upon the current structural state of the cell, *y*. We account for the effect of drugs on the cellular population, then, by taking the product of the cellular apoptosis rate, the drugs’ effectiveness functions, and the respective local drug concentrations *δ_c_p̄*(*t*, *x*)*f̄*(*y*). Multiplying this by the cellular concentration, itself, will yield the degradative sink. As such, the entire source/sink term may be written mathematically as

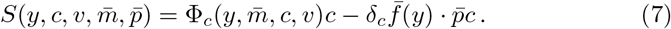

Since, in this particular study, we are interested in the effects of structural heterogeneity on the success of a given cancer population the normalized structural velocity, Ψ(*y*, *m̄*, *p̄*); structurally-dependent growth function, Φ_*c*_(*y*, *m̄*, *c*, *υ*); and the structurally-dependent drug effectiveness function, *f̄* (*y*), are of most interest. Their dependence on structural considerations makes them of particular relevance to the particular situation in which they are applied and so all 3 terms will be specifically defined for the mutational (Section 3) or phenotypic (Section 5) considerations, respectively.

### 2.4. Dynamics in the ECNE, molecular, and drug species

The dynamics of the ECNE, *υ*(*t*, *x*), will be described simply, without spatial dynamics, as growth given by the ECNE remodelling function Φ_*v*_(*c*, *υ*) and the degradation of the ECNE by chemical species. The degradation constant vector will then be given by 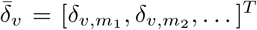 and will have the same number of components as there are chemical species. For any *i*^th^ chemical species that does not degrade the ECNE, the degradation constant *δ_υ_*_,*mi*_ = 0. Our PDE for the ECNE dynamics is then given by

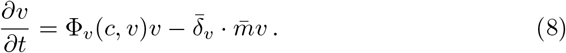

Spatial dynamics of the molecular species vector, *m̄* (*t*, *x*), are given simply by diffusion with its rate vector *D̄* _*m*_. Chemical species are then produced either by the ECNE, and connected network of capillaries, *υ*(*t*, *x*), or the cellular species, *c*(*t*, *x*, *y*), with rates dependent on *y* such that its general expression may be given by the function Φ*̄* _*m*_(*y*, *m̄*, *c*, *υ*). We then assume that environmental factors, which are not directly accounted for, shall contribute to the degradation of molecular species with respective degradation rates of *δ̄_m_*. Dynamics for the molecular species are then collectively written as

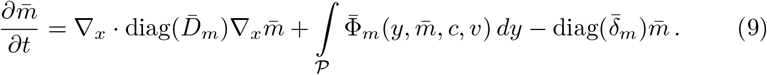

Finally, spatial dynamics for the drug species vector, *p̄*(*t*, *x*), are also given by diffusive dynamics, with a rate vector *D̄* _*p*_. We then represent the input of drug species to the population as a vectorial function, *θ̄*(*t*, *x*), which is to define the drug regimen used by the clinician/scientist in treating the tumour. This will normally be given by a sum of Dirac delta functions centred at the time of injection of the drug but may be given by other forms and will be particular to the experiment that the model attempts to replicate. Finally, we assume that the drug’s effect on the cellular system requires the drug to be taken in by cells and systematically degraded during apoptosis. Therefore, given a drug degradation vector, *δ̄_p_*, this degradation shall be committed by the non-structured cellular population, written as the integral ∫*_𝒫_ cdy*. The complete equation for drug dynamics is then given by

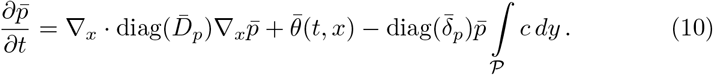

### 2.5. Summary of the General Mathematical Model

We then write the system of PDEs as

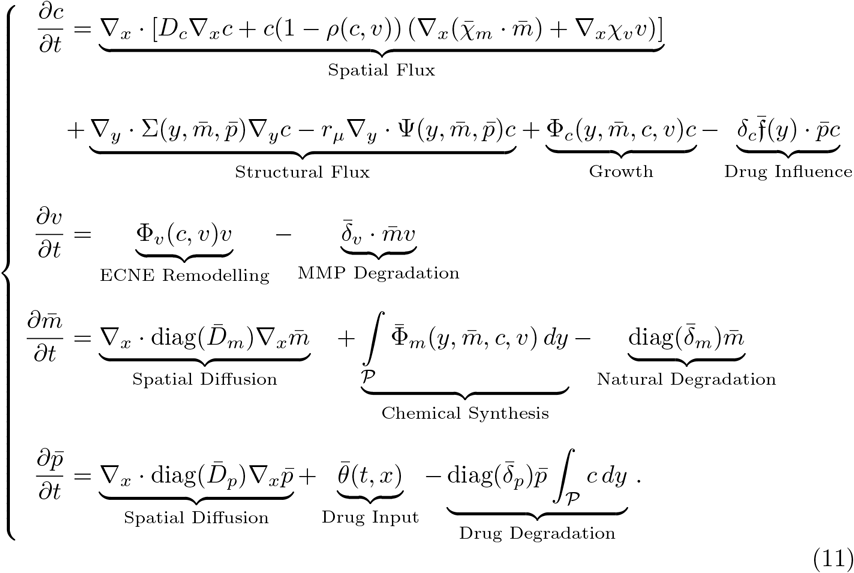

This system of equations (11) is considered together with no-flux boundary conditions in *c*, *m*, and *p̄*. In the case of *c* we consider zero spatial fluxes, and zero structural fluxes on the boundaries of the spatial and structural domains, respectively.

In the following sections, we present an intuitive explanation for the origin of the equations and relations used for two particularisations of this general mathematical system, along with a thorough description of the biological evidence for such phenomena.

## 3. Mutational Evolution and the Establishment of Drug Resistance

### 3.1. Resistance to BRAF inhibitors (BRAFi) and antibody therapies in melanoma

Melanoma is one of the most common cancers of the skin and approximately 50% of melanomas contain a mutation in an oncogene known as BRAF, often appearing at codon 600 [14]. Therefore, BRAF inhibitors (BRAFi) have been the major drug of choice in treating advanced melanoma tumours and their various subtypes. The application of BRAFi as an anti-oncogenic, however, has had mixed results due to the frequent presence of BRAFi resistant phenotypes existing as subspecies within the overall melanoma species [15, 16, 17, 18, 19]. The resistance mechanism could involve activation of collateral signaling path-ways when the main signaling is inhibited [20]. For this reason, simultaneous inhibition of several pathways is often proposed as a possible strategy against resistance [20].

Moreover, recent studies suggest that intravenously injected, water-soluble MAPK activator can overcome, to some extent, the resistance to BRAFi [21]. This, in turn, suggests that the penetration to the inner domain of the tumour is a critical component of the destruction of the resistant cancer cells. Moreover, BRAFi is often used in combination with MEKi in order to target several mechanisms of activation within the MAPK pathway.

In animal models, as well as in patients, relapse occurs systematically several months after treatment with BRAFi [22]. Studies have shown that the adaptations and resistance to BRAFi happen early in the treatment process [23, 24], which may suggest that cancer cells have acquired a resistant state before application of BRAFi.

The order in which drugs are supplied to the tumour may also have a significant effect on the clinical outcome. Progression-free survival rates were higher among those receiving immunotherapy prior to BRAFi than vice versa [25] whereas one particular study looking at treatment with immunotherapy and BRAFi found that preceding BRAFi with immunotherapy does not alter the effectiveness of the drug. Treatment with immunotherapy post-BRAFi, however, gives the patient a particularly poor clinical outcome [26].

One strategy for drug application on the premature tumour has been shown to apparently forestall the resistance to BRAFi. This methodology involved applying the drug to the tumour, for a period of time appearing to demonstrate a reduction in the tumour volume, before removing the drug and repeating the process, again. This method showed mixed results although a significant number of the resistant tumours did not survive the treatment [27].

Tumours that have been shown to have innate BRAFi resistance have further been shown to have increased incidence of mutations in genes known as NRAS [28, 29] and PTEN [30], respectively.

In human liver cells, those cells with an induced PTEN knockdown have been shown to increase the rates of Akt phosphorylation and, importantly, to inhibit Foxo1 signalling [31]. Foxo1, in return, is a transcription factor responsible for mediating the T-cell response to healthy cells [32]. In CD8^+^ T-cells, Foxo1 has been shown to have an intrinsic role in establishing long-lived memory programs that are essential for developing cells capable of immune reactivation during secondary responses to infection [33, 34].

On the other hand, the gene encoding for phosphatidylinositol 3-kinase (PI3K), whose oncogenic pathway is inhibited by PTEN expression, has been shown to reduce the cytokine expression in cells [35], thereby reducing the inflammatory response of the surrounding tissue and limiting T-cell recruitment to the site. Cells with a PTEN deletion might then be protected from immune response through mediation of cytokines and the local apoptosis induced through PI3K/Akt signalling, which may subsequently be overcome by the induction of T-cell hyperactivity induced by ipilimumab – a melanoma-specific immune enhancer therapy.

Therefore, it is possible that melanoma cells undergo sequential genetic alterations in BRAF and PTEN, respectively, and that the pattern in which these mutations occur, along with considerations with respect to competition for nutrients, could explain the build up of resistance to the combined effects of BRAFi and ipilimumab anti-oncogenic treatments.

It may also be that BRAF mutated cells, as a result of causal genomic instability, acquire NRAS mutations which confer resistance. This change, for example, was observed within ovarian cell lines and was predicted to have formed as a result of exon 11 BRAF mutations being insufficient to satisfactorily activate the MAPK pathway, requiring additional NRAS activity [36]. Furthermore, BRAF V600E cells have sufficient MAPK activity such that they do not necessitate supplementary mutation and, as such, display a more positive response to therapy [37], which is supported in the majority of cases of melanoma with a native BRAF mutation [38]. Yet, despite the fact that BRAF and NRAS mutations are described commonly as “mutually exclusive”, NRAS mutations appear in increased numbers of BRAFi resistant tumours [29].

In our model, we interpret the primary and consequent mutation to be that in BRAF and assume, further, that the cell will acquire some further mutation capable of conferring resistance to BRAF inhibitors.

### 3.2. Interpreting the structural dimension for a mutational system

In order to understand how this system of sequential mutations contributes to the cancer cell population’s success at avoiding targeted and immune-enhancement therapies, we must first interpret the structural-, *y*-, dimension. So, letting the cellular population be given by a function *c*(*t*, *x*, *y*) and the ECNE concentrations be given by the function *υ*(*t*, *x*), with *m̄* (*t*, *x*) and *p̄*(*t*, *x*) giving the molecular and drug species, respectively, we observe the bio-mathematical dynamics of such a system in the structure space, *𝒫*.

We also assume, that the cellular species will migrate unidirectionally through the structure space, which is to say that mutations are irreversible. Let the structural mutation variable and space, then, be given by the interval *y ∈ 𝒫*= [0, 1], such that *y* = 0 and *y* = 1 give the extreme states of primary tumour (or as yet without a mutation) and resistant, respectively. For ease, let us also define that *y* = 1/2 defines a BRAF mutation and the state at which the cellular species is most sensitive to BRAFi. Realistically, the ipilimumab immune-enhancer drug will be effective across the entire spectrum of mutations but we assume it to be most effective posterior to BRAF mutation and prior to complete consolidation of resistant features at *y* = 1.

### 3.3. Growth, ECNE remodeling and drug dosing in a mutational system

Let *ρ*(*t*, *x*) be defined as in 2 such that the growth of the cellular species, *c*(*t*, *x*, *y*), shall be dependent upon the unoccupied local volume, 1 − *ρ*(*t*, *x*) and is also dependent upon the nutritional species, *m*_2_(*t*, *x*), being above a given threshold, *θ_m_2__*. The cellular growth rate, with an overall rate parameter *ϕ_c_*, is then written as

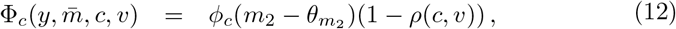

where we consider that growth, in this case, is not dependent upon the mutational status of the cells *y*.

Again, the ECNE remodelling takes place within the unoccupied portion of the local available volume, 1 − *ρ*(*t*, *x*), and with a rate constant *ϕ_υ_*, such that

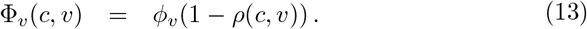

Although we assume here that ECNE remodelling is only dependent on the unoccupied volume, we recognise that more realistically this could depend on fibroblast cells and ultimately on the cell phenotype represented by *y*. Therefore, future iterations of this modelling approach could incorporate more complex remodelling through a redefinition of the Φ_*υ*_ term.

We then endow the system with two molecular species. *m*_1_ is a species that is secreted by the cell species and will act to degrade the ECNE. This can be thought of as a matrix metalloproteinase (MMP) which acts to break-down the ECNE. *m*_2_ is a species which is secreted by the ECNE and acts to the benefit of the cellular species. This chemical species can be thought of as a nutrient or growth factor, the presence of which aids the growth of cellular species.

We further assume that more mutated and aggressive cellular populations will produce MMP molecules at a greater rate, such that their production is proportional to *y*, and that the overall rate constant is given by 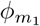. We write this as

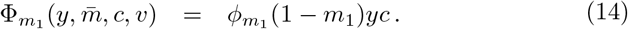

Nutrient, or nutritional species, are produced by the ECNE and with a rate of 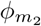, such that

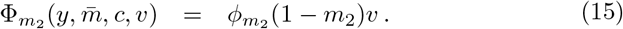

We assume an instantaneous introduction of drug species through the vasculature, which we assume to be proportional to ECNE concentration. The instantaneous nature of this drug introduction mean that we may write this as a Dirac delta function *δ̆*(*t* − *τ*) centered at some time *τ*, whilst its introduction through the vasculature of the ECNE is represented by proportionality to *υ*(*t*, *x*). Then, given that the number of doses of some *j*^th^ drug species, *p_j_*(*t*, *x*), is a natural number, 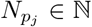, we write that the doses are given at the ordered set of time points 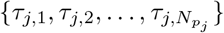, 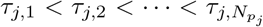. Then the mathematical expression for drug dosing is given by

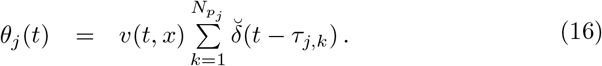

### 3.4. Mutational dynamics in melanoma: Phyletic gradualism or punctuated equilibria?

Patterns in genetic evolution can generally be categorised by the theory of punctuated equilibrium (PE) or phyletic gradualism (PG). PG originates in the theory of Darwinian evolution by natural selection and seeks to explain the variety of species by continuous gradual change [39, 40]. PE, on the other hand, is a currently prominent theory in evolutionary biology that seeks to explain the nature of evolution by natural selection through the prism of large scale genetic and environmental changes, rather than a gradual process [41, 7]. Recent papers in the field of evolutionary biology advance the PE framework as a superior explanation of microbiological, paleontological, and phylogenetic evidence available today [42].

Starting with pioneering contributions of Knudson [43], Cairns [44] and Nowell [45], theory of evolution and population genetics ideas were applied to explain cancer progression. These theories added chromosome instabilities and selection processes to the older idea that cancer results from an accumulation of somatic mutations [46]. Furthermore, the gradual accumulation of mutations over time has been challenged by recent evidence that tumours evolve by a few catastrophic events that generate large scale genome[47] or chromosome lesions[48, 49]. These findings suggest that cancer genomes evolve by PE, being thus able to acquire quickly new capacities such as invasiveness and drug resistance [50, 51, 52]. This PE can be explained on a more microscopic level by assuming that the intermediary stages of mutation, although significant, happen more quickly and to greater effect under certain optimal conditions. Conflictingly, gradualism would convey a sense of regular and linear progression within the phyletic tree of the cancer species with little or no change in the rate of mutation.

Single-cell genetic analysis reveals clonal frequencies and phylogeny patterns of evolving tumours [53, 54, 55, 56]. Various clones have heterogeneous survival properties in the presence of drugs; as a result of this selection pressure, drug resistant clones can become predominant. For instance, mutations of the genes BRAF and NRAS are well known to be driver mutations for melanoma [57, 58, 59, 60]. The wealth of literature on melanomal branching evolution has identified BRAF as the major trunk driver mutation and NRAS or MEK1 as the major branch driver mutations [61, 62]. It has also been recognised that the targeted treatment of genetically evolved melanoma results in a reduction of their heterogeneity [59], as only drug resistant genetic variants survive, but not in their eradication.

For the sake of simplicity, in our model we consider that only two mutations can occur, and that their occurrence is sequential.

### 3.5. A structural flux function in a mutational system

To clarify the mathematical evolution of our cancer cell population, we must more clearly define how the population changes in structure, through the normalized structural velocity Ψ(*y*, *m̄*, *p̄*) (further discussion in Appendix A.1). This function is intended to represent the velocity of any given cell in the *y*-direction (in other words, the mutation rate), for given current structural state (*y*-coordinate) and local nutritional condition, *m*_2_(*t*, *x*). We shall define a separate normalized structural velocity for both a PG and a PE assumption.

In the case of PG, we wish for the evolution of this population to be steady and regular throughout the domain, such that the mutation rate must fundamentally be constant throughout the domain. Then, in order to ensure that our population does not migrate beyond the boundaries of the domain, *y* = 0 or *y* = 1, we set the values of the normalized structural velocity to 0 at these locations, yielding no mutation at these biological positions (Fig. 2a).

**Figure 2:**
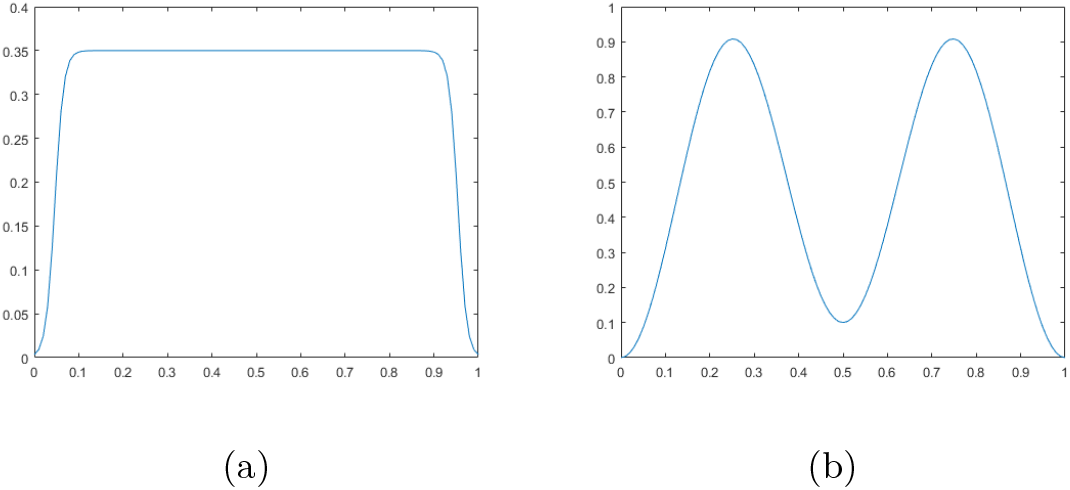
Normalized structural velocity, Ψ(*y*, *m̄*, *p̄*) for the (a) phyletic gradualism (PG) and (b) punctuated equilibrium (PE) assumptions.

In the PE case, we require for the mutation rate to be significantly greater in periods between mutational realisation that at those positions themselves. Therefore, we represent the normalized structural velocity as a bimodal function with velocity maxima positioned between the mutational states. Likewise with the PG function, however, we require for the PE paradigm to yield a 0, non-mutational behaviour at the boundaries of the domain (Fig. 2b). Remember, given that these function represent the rate of mutation, a higher value of Ψ(*y*, *m̄*, *p̄*) will convey a faster rate of mutation whilst a lower value will convey a more quiescent state, where change is somewhat slower.

For the sake of simplicity, we do not consider genetic diversification and structural diffusion in this context.

### 3.6. Drug effectiveness functions in a mutational system

The drug effectiveness is given by a vector valued function *f̄* (*y*):= [*f*_1_(*y*)*, f*_2_(*y*)]*^T^*, where *f_i_*(*y*) gives the effectiveness of its corresponding *i*^th^ drug, *p_i_*(*t*, *x*). For simplicity, we assume that each of these functions is given by a Gaussian function centred at its point of greatest structural significance, or the structural location in *y* at which it is most effective against cancer cells.

Now, since *p*_1_(*t*, *x*) is define to be a BRAFi therapy and we have defined that the BRAF mutation is fully realised at the structural location *y* = 1/2, we assume that *f*_1_(*y*) attains its maximal value at *y* = 1/2 (Fig. 3 *green*). The considerations for ipilimumab are somewhat more numerous and difficult to entirely confirm but are, for our purposes, limited to the following. Firstly, we assume that immune cells should largely ignore healthy cells without a mutation such that there effectiveness at *y* = 0 should be negligible. Moreover, we know that cancer cells will eventually become resistant even to this immune-enhancer therapy and, as such, the value of effectiveness function must be sufficiently low in the neighbourhood of *y* = 1, so as to allow this resistance phenomenon to manifest. Likewise, immune cells require the expression of some protein on the surface of any given cell in order to identify its genetic properties; as such, we assume that only as the BRAF mutation becomes realised, near *y* = 1/2, shall the ipilimumab therapy begin to have a significant effect. Given these considerations, we place the maximum of f_2_ at *y* = 3/4 (Fig. 3 *red*).

**Figure 3:**
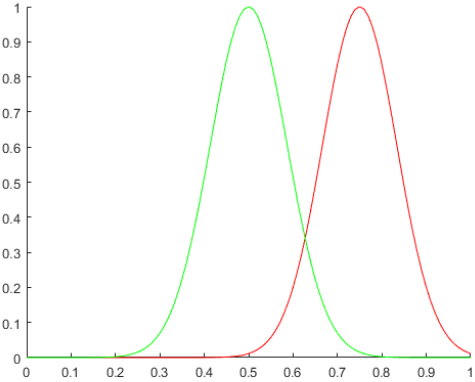
Distribution for the drug effectiveness functions *f*_1_(*y*) (*green*) and *f*_2_(*y*) (*red*).

## 4. Results for the Mutational System

Primarily, in the application of this system to studying the death and regrowth models of tumour resistance in mice, we wished to know whether or not our *in silico* model was able to recapitulate *in vivo* results. In the process of exploring this potential in the model, we attempt to asses the ability of either phyletic gradualistic or punctuated equilibrium assumptions, on the tumour’s evolution, were more able to consistently capture this phenomenon (Section 4.1). Secondly, we wished to test whether, given knowledge of sequenced treatments’ ability to succeed in the ablation of the tumour, we could draw conclusions about the sequencing of treatments and their relative success (Section 4.2). In line with this, we tested periodic treatments to understand what the heterogeneity in initial conditions of the tumour could teach us about the outcomes for treatments (Section 4.3) and, finally, what effect a heterogeneous environment would have on these above conclusions (Section 4.4); whether results would be conserved or altered in the presence of a heterogeneous spatial conditions.

In order to test these scenarios, the *in silico* experimental approach was primarily as so: We began by choosing a melanoma mouse model for which one could attempt to tune our parameters and, effectively, challenge the model. The model that we chose for this task was that of Perna *et al.* who explored the explosive regrowth of tumours after some post-treatment dormancy period [22], amongst other things. Once we had used this *in vivo* model to tune and test our mathematical *in silico* model, we would use other biological models in order to challenge the mathematical model with no further doctoring of the mathematical model or its parameters. For this challenge we chose, initially, that of Thakur *et al.* [27].

Thus, we obtain that these mutations occur at maximal probabilistic rates of approximately 1.9 × 10^−2^ genetic events per day. This corresponds to acquiring a genetic mutation every 40-50 days posterior to some precursor event, where we consider only 2 such events. This is supported by the fact that tumours planted in the mouse species show significant change in expression pattern after 25-45 days [63, 64], where below 40 days BRAFi was a largely successful treatment [65], and mouse models show significant behavioural change in the cancer cell dynamics after 100 days since inocculation [22].

Proliferative and degradative parameters were chosen to be in line with previous models and were fine-tuned for the mouse model considered, based on tumour growth rates observed in tuning experiments [22]. All of these values are summarised in Table B.1.

### 4.1. Punctuated equilibrium (PE) assumptions are more consistent with in vivo experimental results than phyletic gradualism (PG) assumptions

Given certain initial conditions for the cellular population, namely an initial structural distribution centred at *η* = 1/20, both PE and PG assumptions can give rise to the characteristic death and regrowth curves, albeit with differing characteristics (Fig. 4). In both cases, one observes an initial growth phase which is quickly stunted and violently reversed by the introduction of the drug species at *t* = 45. This is followed by a period of dormancy or ‘tolerance’ before the characteristic resistant growth (or regrowth) phase, which is of particular interest to our current study. Observe, initially that the regrowth phase manifests a an earlier time point and with a faster growth rate under PE assumptions than with PG assumptions.

**Figure 4:**
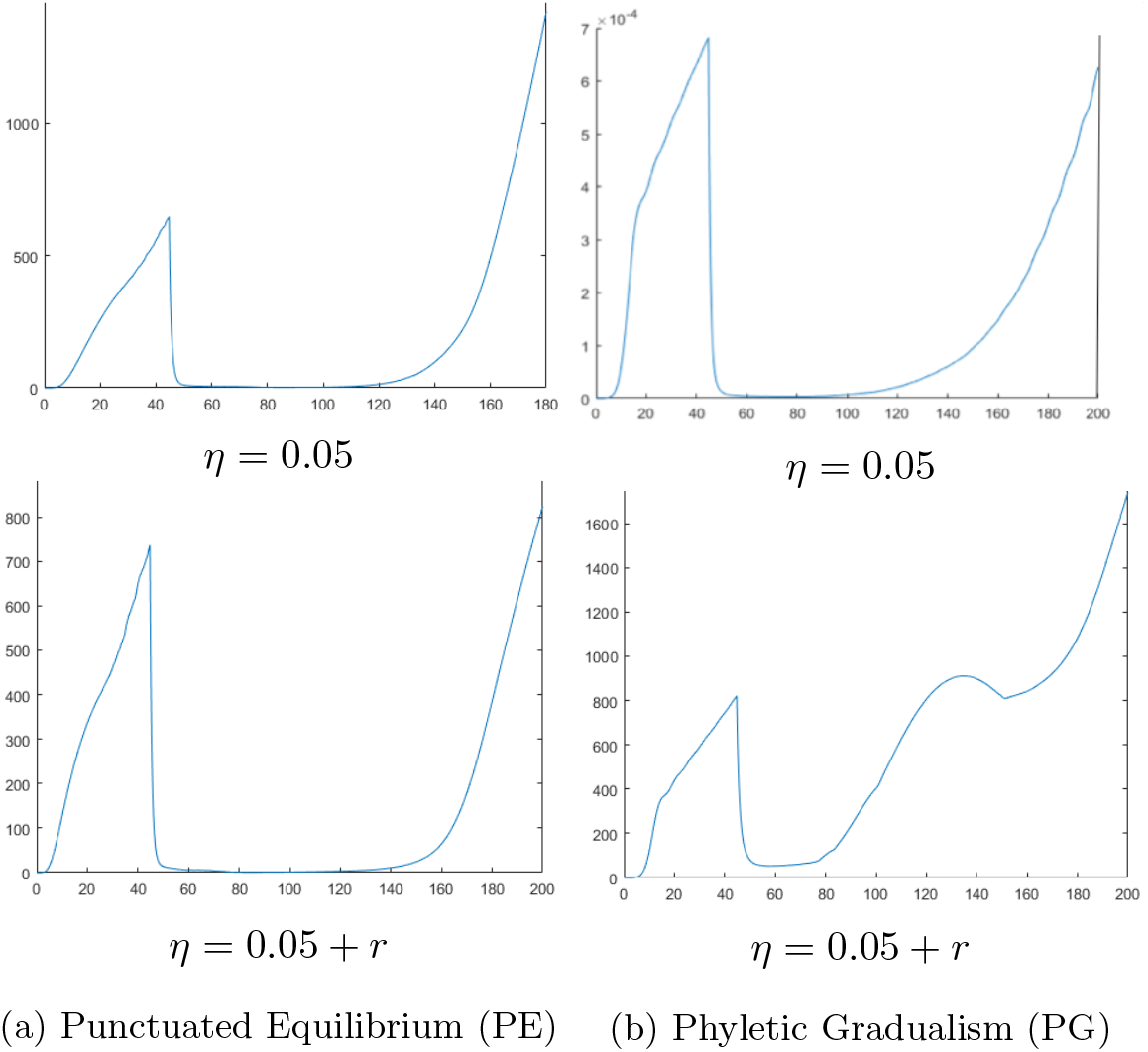
Punctuated evolution is more consistent with biological results than gradual evolution. Tumour volume graphs for (a) punctuated equilibrium (PE) and (b) phyletic gradualism (PG) assumptions under a simply BRAFi therapy option applied at *t* = 40 for initial conditions of *η* = 0.05 or *η* = 0.05 + *r*, where *η* represents the initial mean location of the tumour cells along the phenotypic dimension, and *r* = 0.1 represents a perturbation.

Now, observe that inducing a significant (200%) perturbation in only the position of the initial conditions, we evoke dramatically differing behaviours from our two *in silico* tumours (Fig. 4). For the case of PE, the rate at which our tumour regrows to its pre-treatment volume is much slower but the death and prolonged dormancy phases are conserved between these two experiments (Fig. 4a). Under the assumptions of PG, however, one observes at all time points a tumour volume with a significant positive minimum value (Fig. 4b). This shift in the volumes of tolerant tumours to be visible for all time points is not consistent with the results of comparative *in vivo* experiments [22] and, thusly, the initial conditions of a PG model would have to be strictly constrained to some smaller subset of possible conditions in order to maintain its relevance.

In biological, and especially in the case of *in vivo*, experimentation, however, the initial conditions of a given tumour or its new environment may never be strictly limited. This would suggest, due to its robustness to fluctuations in initial conditions, that the PE modelling assumption is most consistent with the results of murine experimentation, since the characteristic death and regrowth curve is conserved.

### 4.2. Sequencing and order of treatments are vital to their success

In order to test the importance of the order of drug treatments on the resistance phenomenon we have first used homogeneous initial conditions for the ECNE. These conditions also preserves the spherical symmetry of the tumour when drugs are applied uniformly on the periphery. Heterogeneous initial conditions leading to non-spherically symmetric tumours will be tested in Section 4.4.

With that understood, in all cases and treatment scenarios the tumours initially respond to treatment, exhibiting a significant period of apoptotic degradation (Fig. 5). Experiments wherein only one treatment was used (Fig. 5a & 5b) show dramatically differing clinical treatment profiles. BRAFi treatment shows an extremely promising tumour response with almost complete erradication occuring within days of treatment but followed by an exaggerated regrowth (Fig. 5a), as seen in murine experiments. Ipilimumab therapy does not show as successful an eradication pattern at earlier time points but is more consistent in quelling its resistance and resulting regrowth (Fig. 5b), although ultimately unsuccessful in eradicating the tumour.

**Figure 5:**
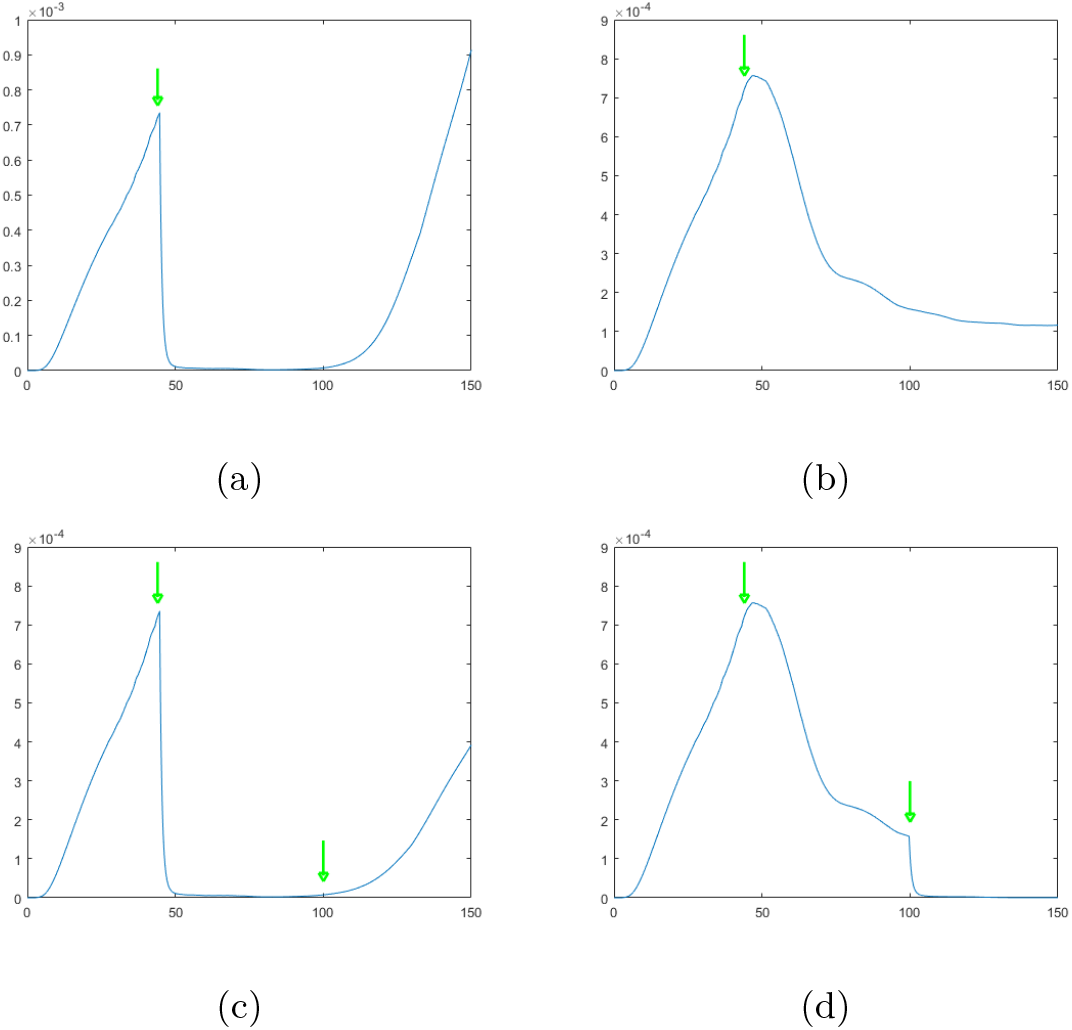
The sequencing of treatments is crucial to success. Overall tumour volume over time, calculated using (B.5), with the drug strategies (a) BRAFi, (b) ipilimumab, (c) BRAFi followed by ipilimumab, and (d) ipilimumab followed by BRAFi; where the drugs are applied constantly after some *t* = 45 (1*^st^ green arrow*) and then *t* = 100 (2*^nd^ green arrow*) when applicable

Observing the therapeutic strategy of utilising a BRAFi treatment followed by an ipilimumab post-treatment is ineffective at destroying the tumour (Fig. 5d). Although the ipilimumab post-treatment is slowing the growth of the now aggressive tumour, it may already be resistant to immunological therapies. The ipilimumab treatment followed by BRAFi post-treatment, however, appears to be extremely effective (Fig. 5d), with a negative growth rate for the tumour volume maintained as of *t* = 1000 (Results not shown). This counterintuitive result may be explained as follows: Firstly, BRAFi appears extremely effective at depleting the tumour volume but is incapable of preventing the resistant escape of subpopulations to higher values of *y* (Fig. 5a). On the other hand, ipilimumab’s effectiveness function is centred at a greater value of *y* than BRAFi’s, making ipilumumab appear less effective but allowing ipilimumab to effectively confine surviving tumour cells at lower values of *y*, where BRAFi remains effective. Therefore, these results would suggest that BRAFi should be used to destroy the tumour once its tendency towards resistance has been stemmed through ipilimumab’s immunological mechanisms.

### 4.3. Oscillatory tumour volumes as a result of periodic treatments do not necessarily imply re-sensitisation

Our second experimental approach was to attempt the experiment of Thakur *et al.* [27] who implemented a periodic treatment regimen for their *in vivo* tumours. This periodic treatments managed to eradicate the death and rapid regrowth phases of those previous experiments and instead resulted in oscillatory dynamics in the tumour volume. Across several cycles of these treatments, some tumours managed to outgrow the drugs and became resistant, although more slowly, whilst others appeared to reduce their volume even over far longer time-periods. The research team explained this by suggesting that the application of less severe treatment regimes may delay the resistance to treatment in solid tumours by failing to encourage the development of such resistance.

Likewise, in our experiments we observed an oscillatory dynamics resulting from the periodic application of smaller dosages to the tumour and subsequent removal of the dose. We found that as we increased the number of independent starting *y* positions in the initial conditions for our cancer cell population, our results gave a greater qualitative agreement with those of Thakur *et al.* [27]. Moreover, we found that there was a strong correlation between the average *y*-position of the initial condition and the final tumour volume at *t* = 160.

**Figure 6:**
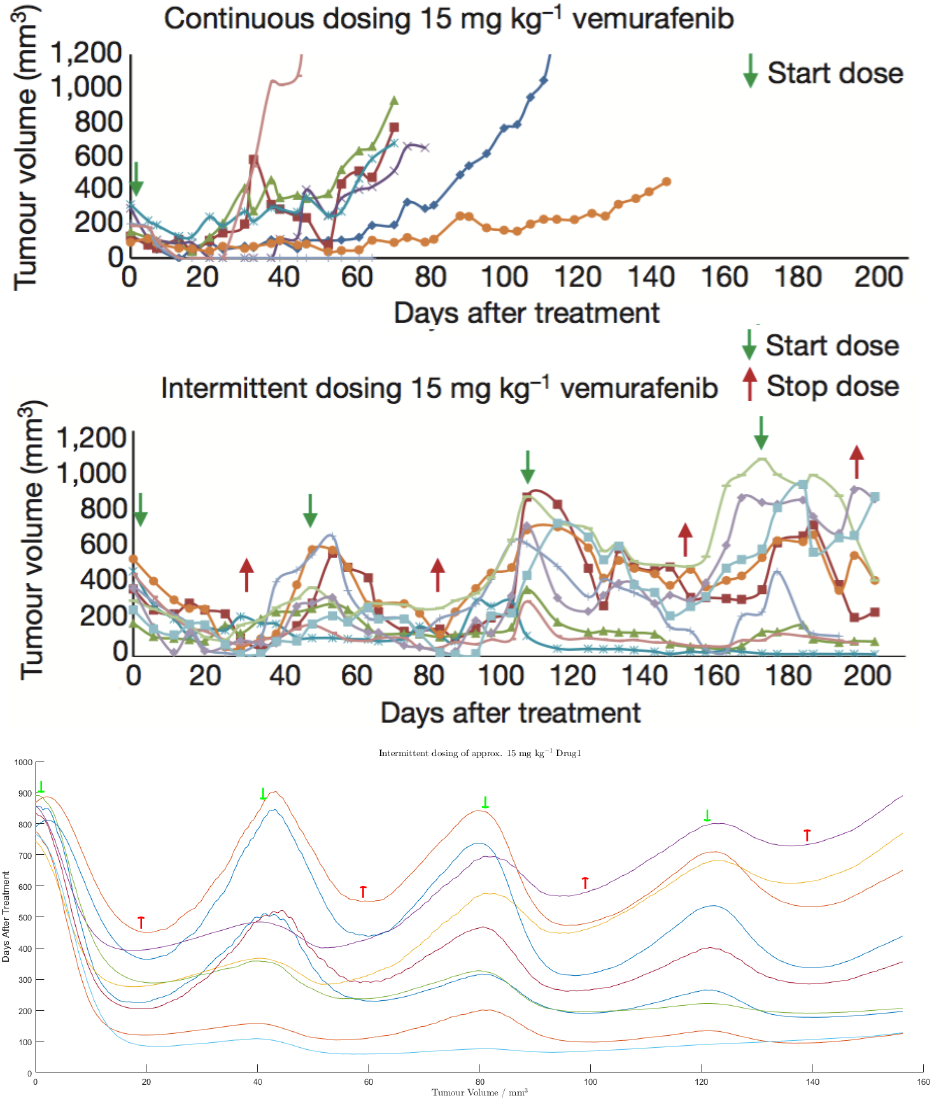
Oscillatory tumour volumes can emerge in the absence of re-sensitisation. The *top* and *middle* panels are figures from Das Thakur *et al.* [27] for *in vivo* melanoma tumours under an intermittent dosing strategy and the *bottom* panel gives the *in silico* results of the same experiments run using the mutational mathematical model. (Licenses applied for from Nature Publishing Ltd.)

These results allowed us to reinterpret this oscillatory behaviour. In our *in silico* model, the acquisition of resistance is certainly not delayed because cells are progressing irreversibly in the *y* direction. In fact, what may be occurring is that in a situation where some number of cells are resistant whilst other are not, these two heterogeneous subpopulations will have to compete for available nutrients in the environment. Not only this but, together, they will consume more nutrients, leaving fewer such nutrients for the resistant subpopulation and leaving a greater subpopulation sensitive to existing treatment options. A dynamical state will be reached where the two sub-populations are oscillating while keeping their volumes bounded.

### 4.4. Drug success rates decay under heterogeneous spatio-environmental assumptions

In order to examine the effect that spatial heterogeneity of the ECNE concentrations and, thusly, the resulting cancer cell population on the longer term effectiveness of targeted and immunological treatments, we considered only that treatment protocol which proved effective in the homogeneous case; namely that of an ipilimumab treatment followed by BRAFi post-treatment. The introduction of spatial heterogeneity whilst maintaining all other factors, in their entirety, was sufficient to cause the degeneration of treatment success into the characteristic death and regrowth curves seen previously (Results not shown, although they may be inferred from figures 7 & 8 *middle*).

**Figure 7:**
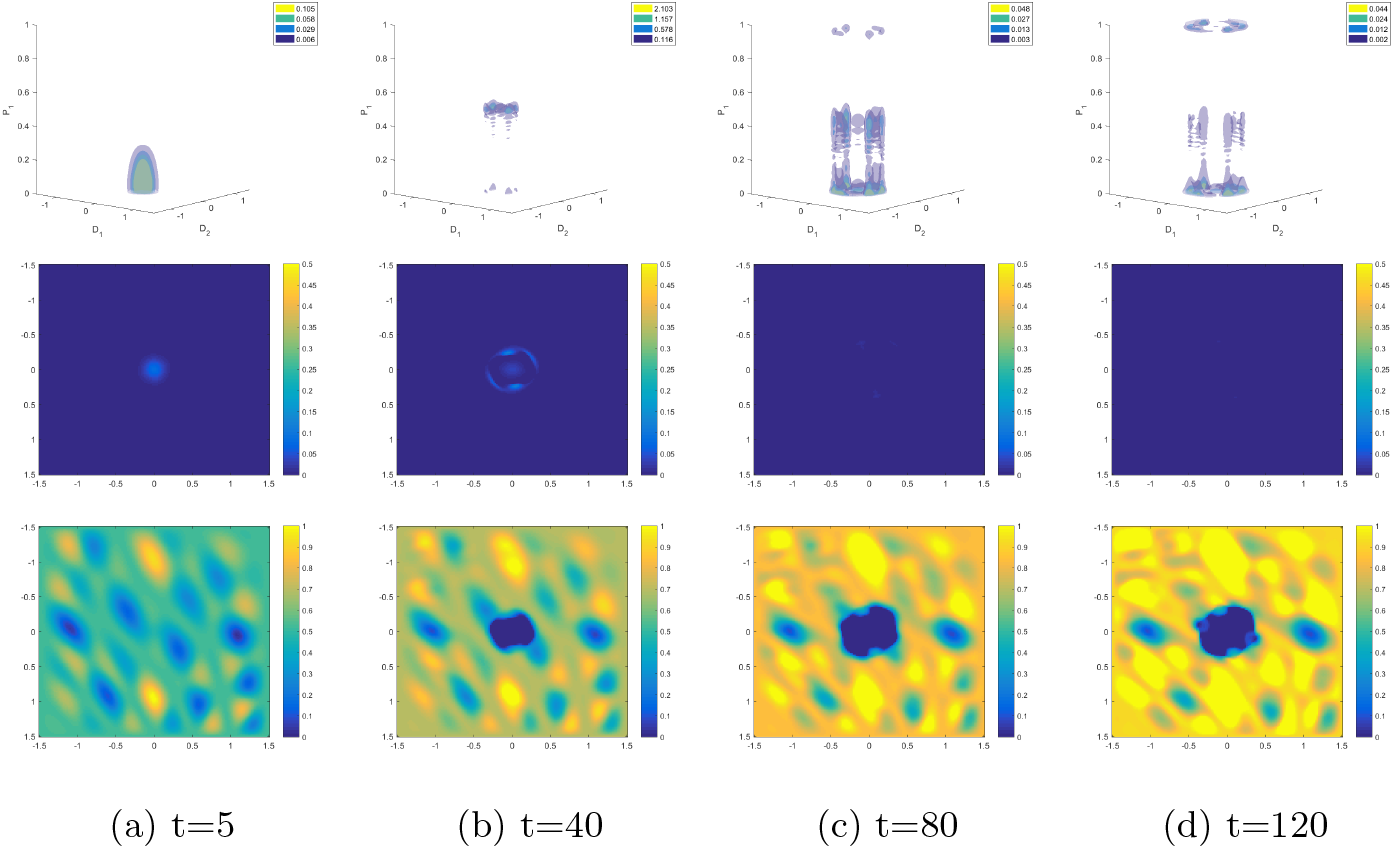
Spatial heterogeneity eradicates treatment success. Panels displaying (*top*) the structured cellular population with space across the lower plane and mutational state given along the vertical axis; (*middle*) the spatial cellular distribution; and (*bottom*) the ECNE density, where ipilimumab treatment is given at *t* = 40 and BRAFi treatment is given at *t* = 100, for time points *t ∈* {5, 40, 80, 120} are shown.

**Figure 8:**
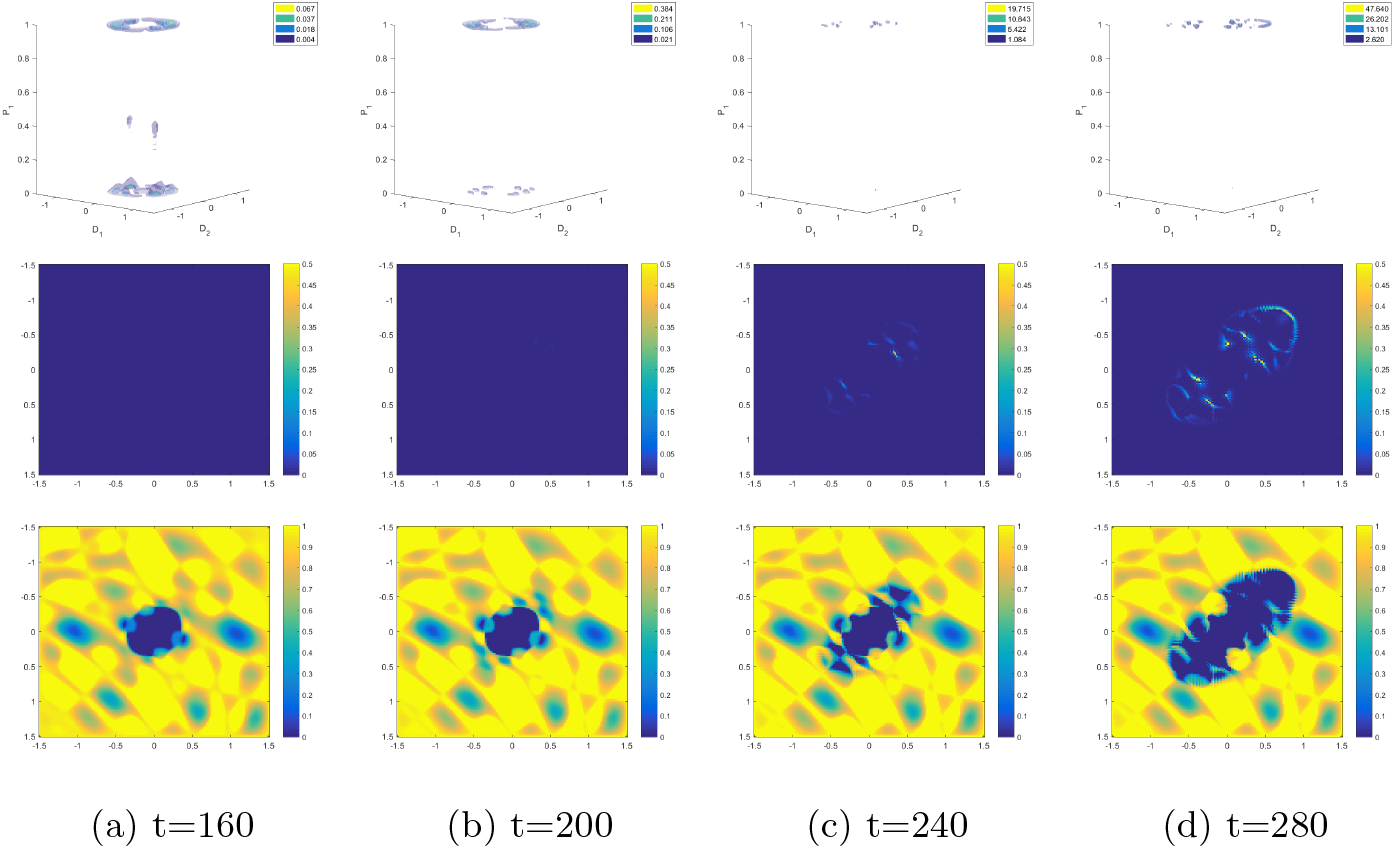
Spatial heterogeneity eradicates treatment success. Panels displaying (*top*) the structured cellular population with space across the lower plane and mutational state given along the vertical axis; (*middle*) the spatial cellular distribution; and (*bottom*) the ECNE density, where ipilimumab treatment is given at *t* = 40 and BRAFi treatment is given at *t* = 100, for time points *t ∈* {160, 200, 240, 280} are shown.

Notice, firstly, that the spatial cancer cell population (Fig. 7 & 8 *middle*) initially spreads to the nearby regions of elevated ECNE concentration, prior to treatment. As the treatment is applied, and the regions of highest cell population coincide with the regions of highest ipilimumab concentration, the cell population is reduced to invisibility for some times 40 < *t* < 200. It should be understood, here, that under a great evolutionary selective pressure only very few cells survive these initial waves of treatment but those cells which do survive will be completely resistant to both treatments. At this time, and with almost the entirety of the surviving cellular population being resistant to both BRAFi and ipilimumab, the cellular population begins to regrow at regions of highest nutritional content, or ECNE.

One may observe this dynamic in the spatio-structural cellular population, progressively over the entire time domain. Consistently with the punctuated equilibrium assumptions within the model, one notices a pulsatile movement of the cellular subpopulations between *y* = 0 and *y* = 1/2, and again towards *y* = 1 (Fig. 7 & 8 *top*). In particular, however, the first time at which the cancer cell population has been visibly eradicated (Fig. 7c), the visible coincidence of those areas of low ECNE concentration with those cancer cell clusters at the most elevated value of *y*. In other words, the difference in the heterogeneous case, as compared with the homogeneous case, is that the cancer cell population is able to preferentially avoid drug-induced apoptosis by remaining in regions of low ECNE and drug concentrations, which allows the cellular population to become resistant before migrating to regions of high nutrition and increasing their collective proliferation rate.

This demonstrates that particular prudence must be paid during consideration of spatial factors in the study of drug resistance and strategy. One should also notice the clinically difficult tumour that results from this method of treatment (Fig. 8d *middle*) and the nature of the underlying environmental infrastructure, or ECNE. The tumour is viable although sparsely populated which raises significant questions about the ability to remove such a tumour, surgically. The approach to treating such a patient would classically be to use chemical means, which have now been exhausted and given rise to a uniformly resistant tumour.

## 5. Metabolic Remodeling and the Re-Establishment of Drug Sensitivity

Recent studies have looked at the effect of BRAFi on the human melanoma PDX lines implanted in the immunodeficient mouse and found that this drug is largely ineffective, implicating a role for the immune system in its functioning. This result is contrasted with the effectiveness at eradicating the tumour with BRAFi+MEKi, again with the characteristic relapse curve [66].

These same studies have suggested that after a primary phase of treatment, and subsequent washing of the drug species from the tumour, the cancerous cells may regain their sensitivity [66]. This is illustrated in the cells’ recapitulation to later phases of treatment and suggests that some metabolic, or other, plasticity may lead to the observed resistance to BRAFi and MEKi. This plastic response may be reversed upon the removal of the drug and is believed to be as a direct result of stress on the cells themselves.

Beyond these conclusions of the study, the observation is made that the system remains the genetic equal of the precursor tumour at every stage during this adaptive process. This suggests that a population-wise phenotypic switch occurs from populations that are composed 1% of epigenetically resistant cells, prior to treatment, to being comprised 70% of this cell type, post-treatment and post relapse [66]. Little is known about the phenotypic status of the tumour immediately prior to the secondary round of BRAFi+MEKi dosing.

Moreover, cutaneous tissue is naturally and significantly heterogeneous in its composition and, being the tissue furthest from the major vasculature, is greatly dependent on the arterial supply of oxygen and other nutritional components of the cellular system. In areas with the lowest such supplies of oxygen, cells switch their metabolism from mostly oxidative phosphorylation (oxphos) to glycolysis. Using, then, BRAFi and MEKi in order to inhibit the glycolytic pathway [67, 68] induces an excessive stress regimen within the cell. It has been suggested that, under such powerful metabolic stresses, the cell will diversify its metabolic behaviour in order to attempt an increase in efficiency. This switching between glycolytic and oxphos modes of metabolism may, therefore, be instrumental in facilitating the avoidance of targeted inhibition within cancer cells; cancer cells may use oxphos metabolism to avoid the targeted inhibition of glycolysis [69].

This, however, implies that we are now existing within a different paradigm with respect to the evolution of the cells in response to drug application or, perhaps, in general. To begin with, we recall that *p*_1_(*t*, *x*) is given by the spatio-temporal concentration of BRAFi and we, now, redefine that *p*_2_(*t*, *x*) should be given by the spatio-temporal concentration of MEKi, a second metabolic inhibitor of glycolysis.

### 5.1. Re-interpreting the structural dimension for a metabolic system

In order to capture the re-sensitisation phenomenon, we must reinterpret the structural *y* variable to take into account the newfound plasticity of the cellular population. We assume that the effect of the drugs and the variability in the cellular population may be adequately illustrated through the cellular pathways involved in the metabolism of glucose; namely those of glycolysis and of oxphos. Given that a given glucose molecule, may be metabolised through the utilisation of either one of these pathways, but not both, we may represent the structure of the cell as the proportion of glucose sent to glycolytic pathways as opposed to oxphos pathways; such that *y* = 0 represents 100% of glucose being metabolised through glycolysis, and 0% by oxphos, whilst *y* = 1 represents 0% of glucose being metabolised through glycolysis, and 100% by oxphos (Fig. 9).

**Figure 9:**
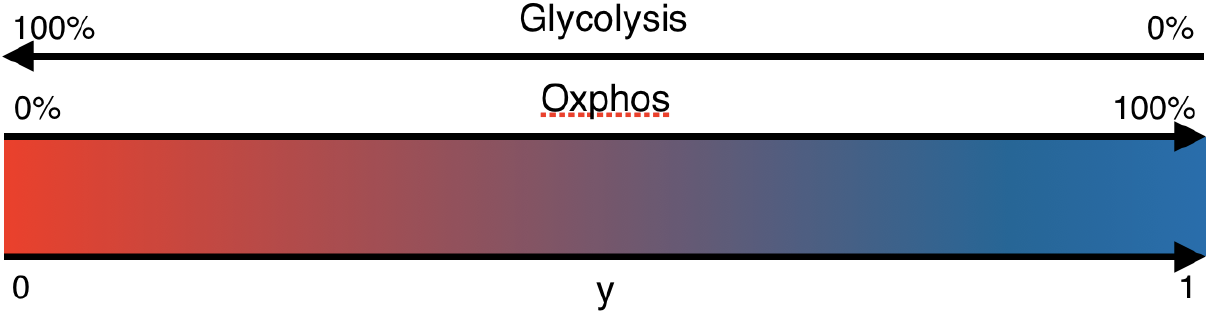
Visual reinterpretation of the structural *y* variable to account for the metabolism of glucose molecules proportionally and competitively through glycolytic and oxphos pathways, respectively.

### 5.2. A cellular growth function in a metabolic system

Likewise with our previous paradigm, we assume that proliferation requires the presence of nutrients, *m*_2_(*t*, *x*), above a certain threshold, 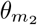. As was recognised by Warburg in 1956 [70], and was subsequently termed the Warburg effect, highly proliferative cancer cells appear to preferentially utilise glycolytic pathways to synthesise membrane lipids and other essential components from glucose. Therefore, we assume that there exists some underlying proliferation rate, *ϕ_c_*_,1_, which is common amongst all cells and a further ‘Warburg’ proliferation rate, *ϕ_c_*_,2_, which is contributed dependent upon the degree to which the cell utilises glycolysis; as the cell utilises the glycolytic pathways to a greater extent, its proliferation rate shall increase concurrently. Moreover, since we are particularly interested in the cell’s ability to absorb and utilise available nutrients in the environment, we modify our competition assumptions so that the cellular population’s proliferation will not be inhibited by the presence of the ECNE but will rather simply increase the pressure on the ECNE itself. Thusly, we replace the unoccupied volume term by 1 − ∫*_𝒫_ c dy* and write the full growth term as

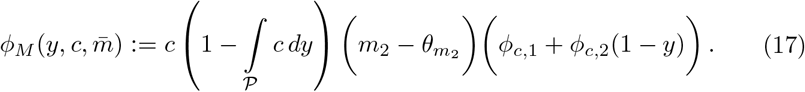

### 5.3. A structural flux function in a metabolic system

The cell is biologically engineered to complete its cell cycle and evolution has selected for cellular populations who are particularly efficient at achieving this goal. Therefore, given that a cell requires nutrition and the ability to freely adapt in order to achieve this objective, if the cell is deprived of its essential environment then it will take extreme measures in order to continue to proliferate. We here define stress, or ‘stressed conditions’, as those conditions which are not conducive to cellular metabolism and proliferation. In particular, those scenarios which would lead the cell to feel ‘stressed’ are given explicitly by nutritional deprivation or targeted inhibition of metabolically essential genes, such as BRAF or MEK. Therefore, we define the weighted stress term as 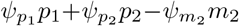, where 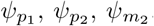, are positive weights such that 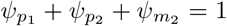. Under stress, the cell shall randomly diversify its behaviour; each cell becoming stochastically more or less oriented towards glycolytic metabolism such that the population, as a whole, becomes more metabolically diverse. Therefore, we may represent this at the population level by a structurally diffusive behaviour. The structural diffusion coefficient Σ(*y*, *m̄*, *p̄*) is proportional to the weighted stress, therefore

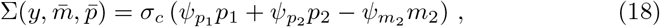

where *σ_c_* is a positive constant. In the absence of stress, the cell population relaxes by advection to the preferential metabolic state *y* = *ω_c_*. The relaxation rate is proportional to the weighed non-stressed factor defined as 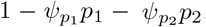. Thus, the normalized structural velocity reads

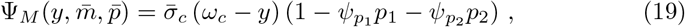

where *σ̄_c_* is a positive constant.

### 5.4. Drug effectiveness functions in a metabolic system

The drug effectiveness functions for BRAFi and MEKi, *p*_1_(*t*, *x*) and *p*_2_(*t*, *x*) respectively (further discussion in Appendix A.2), are given simply by Gaussian functions centred at 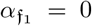 and 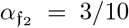 respectively. We write these mathematically

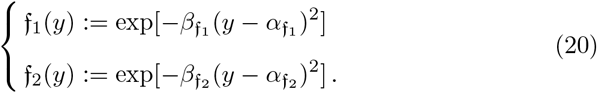

whilst the widths of these Gaussian functions are uniform with 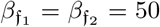 (Fig. 10), in order to replicate results from the murine models from [66].

**Figure 10:**
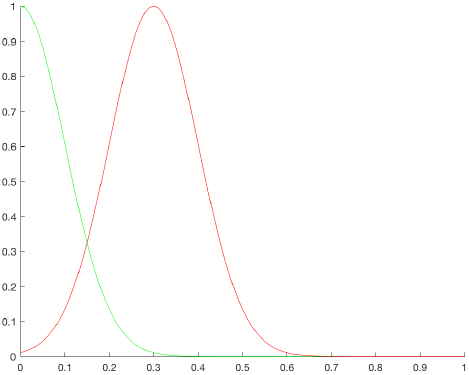
Distributions for the drug effectiveness functions *f*_1_(*y*) (*green*) and *f*_2_(*y*) (*red*).

## 6. Results for the Metabolic System

Again, our primary motivating factor for these metabolically plastic systems was to understand whether these mathematical models were capable of recreating or predicting the complex dynamics underlying *in vivo* results (Section 6.1). Beyond this, we wished to try to understand the spatio-metabolic dynamics of the tumour which are allowing resistance to develop (Section 6.2). Finally, given the complexity of the plastic model, we wished to know what the dynamics of the cellular population, under the influence of drugs, might tell us about the reaction of this population to treatments and the clinical significance of this reaction (Section 6.3).

In order to test this *in silico* model, we attempted to recreate the conditions in the experiments run by Rambow *et al.* [66]. In these experiments, mice were given a PDX melanoma and the tumour was allowed to grow for some initial period without treatment. Tumours were then treated with BRAFi+MEKi combination therapy at a time point which corresponded to 80 days of growth (*t* = 80) in our *in silico* tumours. As the tumour developed resistance to the treatment, the dose was released at the time point corresponding to the volume of tumour increasing to approximately 50% of its volume prior to treatment, which we selected as *t* = 210 in our tumours. A final dose was given after approximately 30 days of unimpeded growth, at *t* = 240.

### 6.1. Resistance and re-sensitisation dynamics are captured by plastic, metabolic in silico modelling

As is the case with the *in vivo* experiments, we observe the death, tolerance, and regrowth pattern within the tumour (Fig. 11). This is then followed by a period of rapid, unimpeded growth due to the removal of drugs from the tumour. It is important to notice that upon the second wave of treatment, the tumour is again eradicated entirely for some brief period before becoming resistant more rapidly on this second occasion (Fig. 11). This correlates qualitatively with the *in vivo* results but may not be explained by a mutational model since those resistant cells would not reestablish their sensitivity to treatment. This effect is termed ‘re-sensitisation’ and may be biologically and clinically significant.

**Figure 11:**
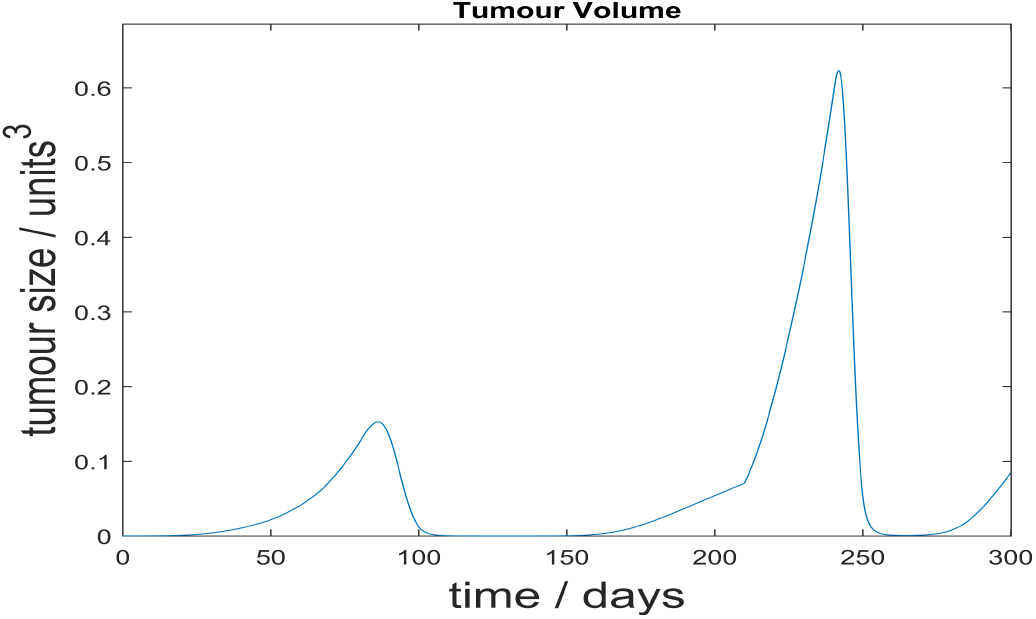
Resistance and re-sensitisation dynamics are captured by *in silico* modelling. Graph displaying the tumour volume of the metabolic tumour model over the duration of the *in silico* experiment, with continuous doses given from *t ∈* {80, 240} and a drug holiday initiated at *t* = 210

In order to more accurately capture the results of the biological, experimental approach we use a lower dosing rate in this model. Also, the dose was applied uniformly in time between the start and the end of the treatment, instead of instantaneously (we used Heaviside functions instead of Dirac functions for the drug temporal profiles). This ensured a more gradual switch from the initial growth stage in the tumour to a drug-sensitive apoptotic phase, prior to tolerance (Fig. 11). Moreover, the primary regrowth stage appears to be damped in comparison to the mutational model under BRAFi treatment, alone, but this could be explained by the supplementary dosing of the tumour with MEKi, stunting regrowth to a greater extent.

### 6.2. Temporary oxphos metabolism may allow cancers to evade targeted treatments

Recall that lower values in *y* are associated with more glycolytic modes of metabolism, where higher values of *y* are associated with more oxphos modes of metabolism and that each of these structural *y*-coordinates is associated with a 2D spatial *x*-coordinate. Moreover, a *green* encircled 1 in the upper right-hand corner of a graphic shall signify that the tumour is under BRAFi treatment, where a *red* encircled 2 in the upper right-hand corner of a graphic shall signify that the tumour is under MEKi treatment (Fig. 12–15).

**Figure 12:**
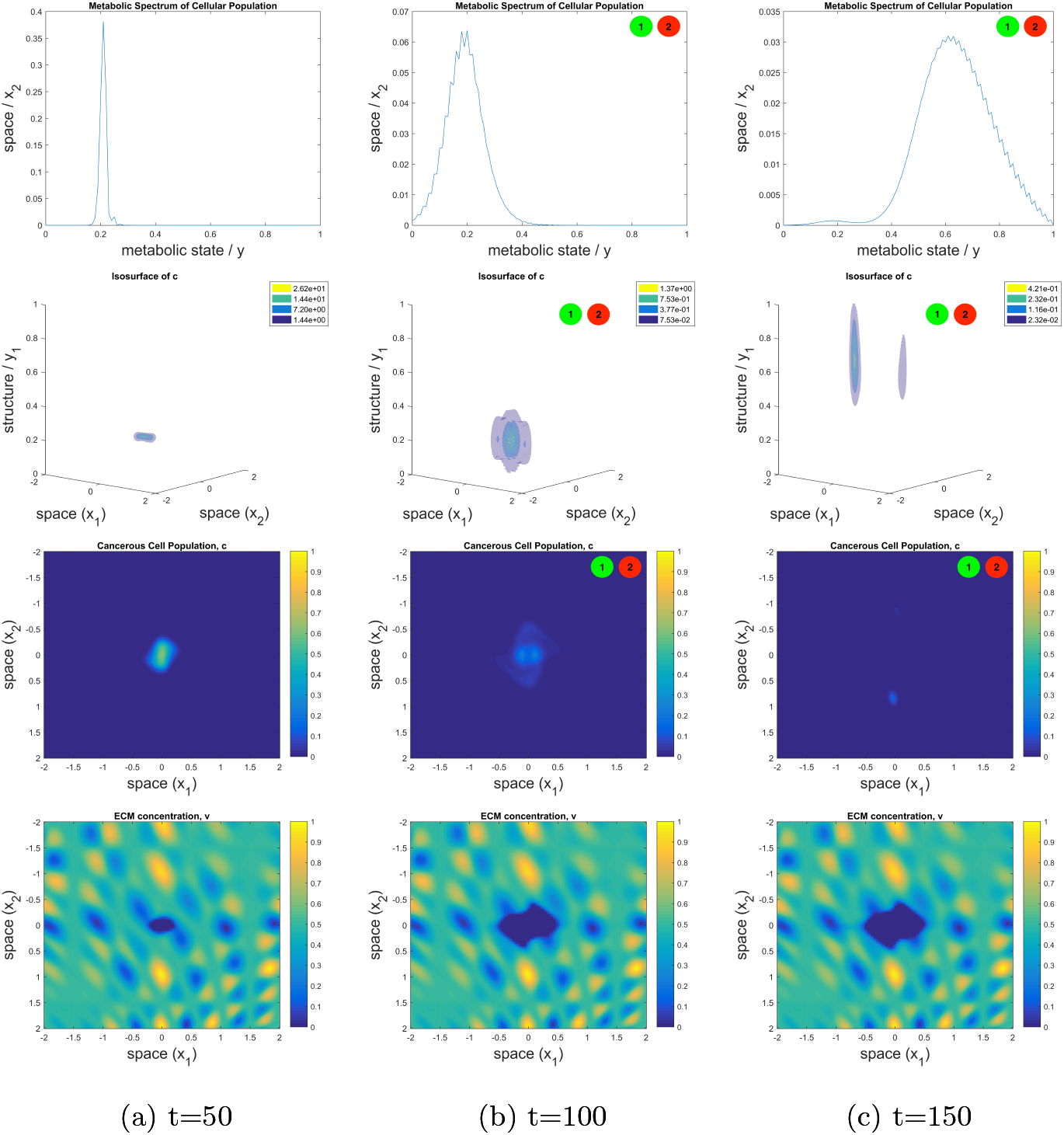
Tumours use oxphos metabolic pathways to resist targeted inhibition of glycolytic pathways by BRAFi and MEKi therapies. Shown are the phenotypic distribution (1*^st^ row*); the spatio-phenotypic surface distributions (2*^nd^ row*); and spatial distribution (3*^rd^ row*) of the cellular population. The spatial distribution of the ECNE, with colour-bar, is also shown (4*^th^ row*), for completeness and in order that one can place the tumour within its environmental context. All figures are given at times *t ∈* {50, 100, 150} within subfigures (a), (b), and (c) respectively. Within the surface plots, the colours represent surfaces of approximately equal concentrations within the spatio-phenotypic context of the cell gradiated from lowest to highest concentration as purple, blue, green, then yellow.

**Figure 13:**
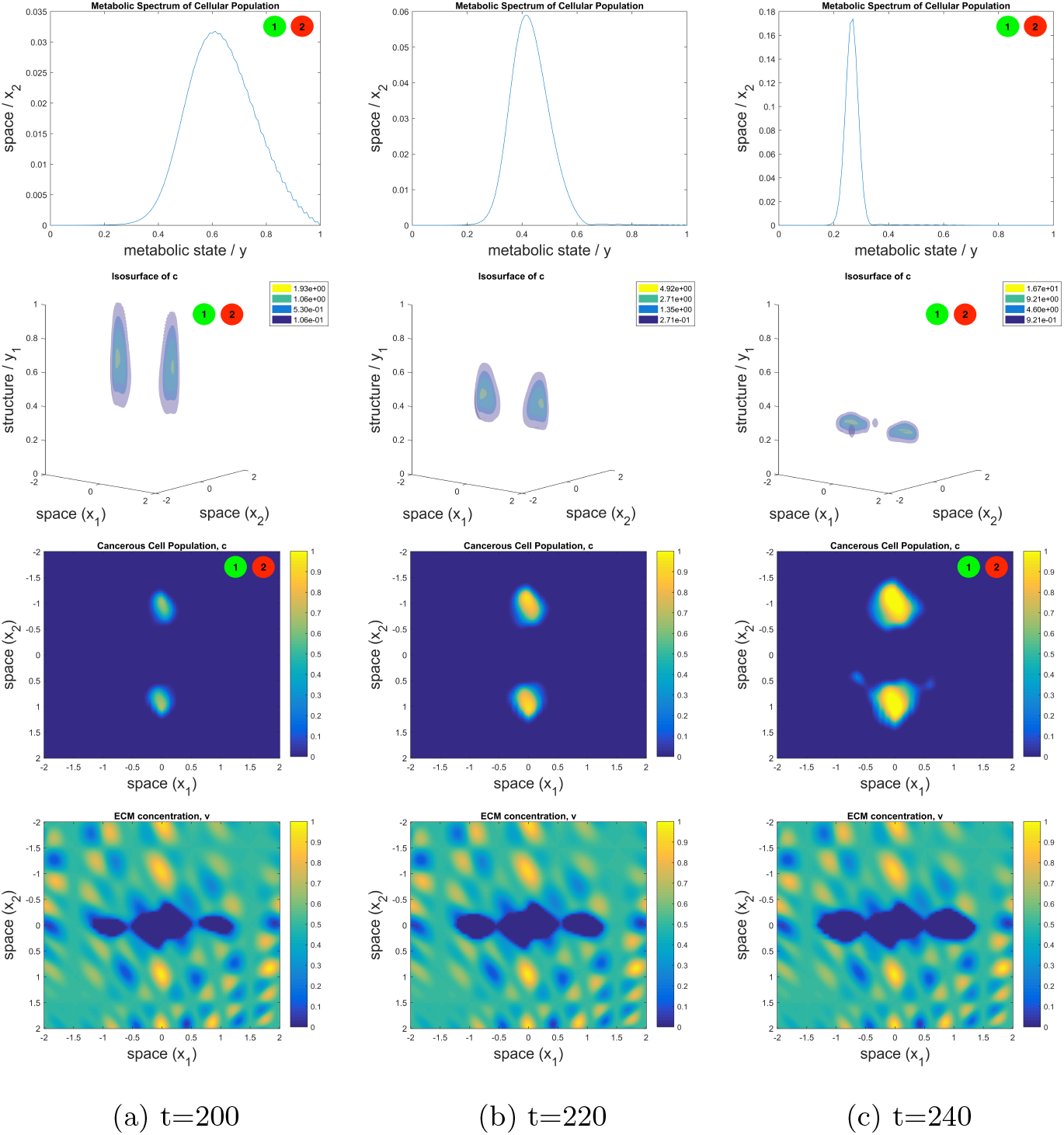
Tumours use oxphos metabolic pathways to resist targeted inhibition of glycolytic pathways by BRAFi and MEKi therapies. Shown are the phenotypic distribution (*1^st^ row*); the spatio-phenotypic surface distributions (*2^nd^ row*); and spatial distribution (*3^rd^ row*) of the cellular population. The spatial distribution of the ECNE, with colour-bar, is also shown (*4^th^ row*), for completeness and in order that one can place the tumour within its environmental context. All figures are given at times *t ∈* {200, 220, 240} within subfigures (a), (b), and (c) respectively. Within the surface plots, the colours represent surfaces of approximately equal concentrations within the spatio-phenotypic context of the cell gradiated from lowest to highest concentration as purple, blue, green, then yellow.

**Figure 14:**
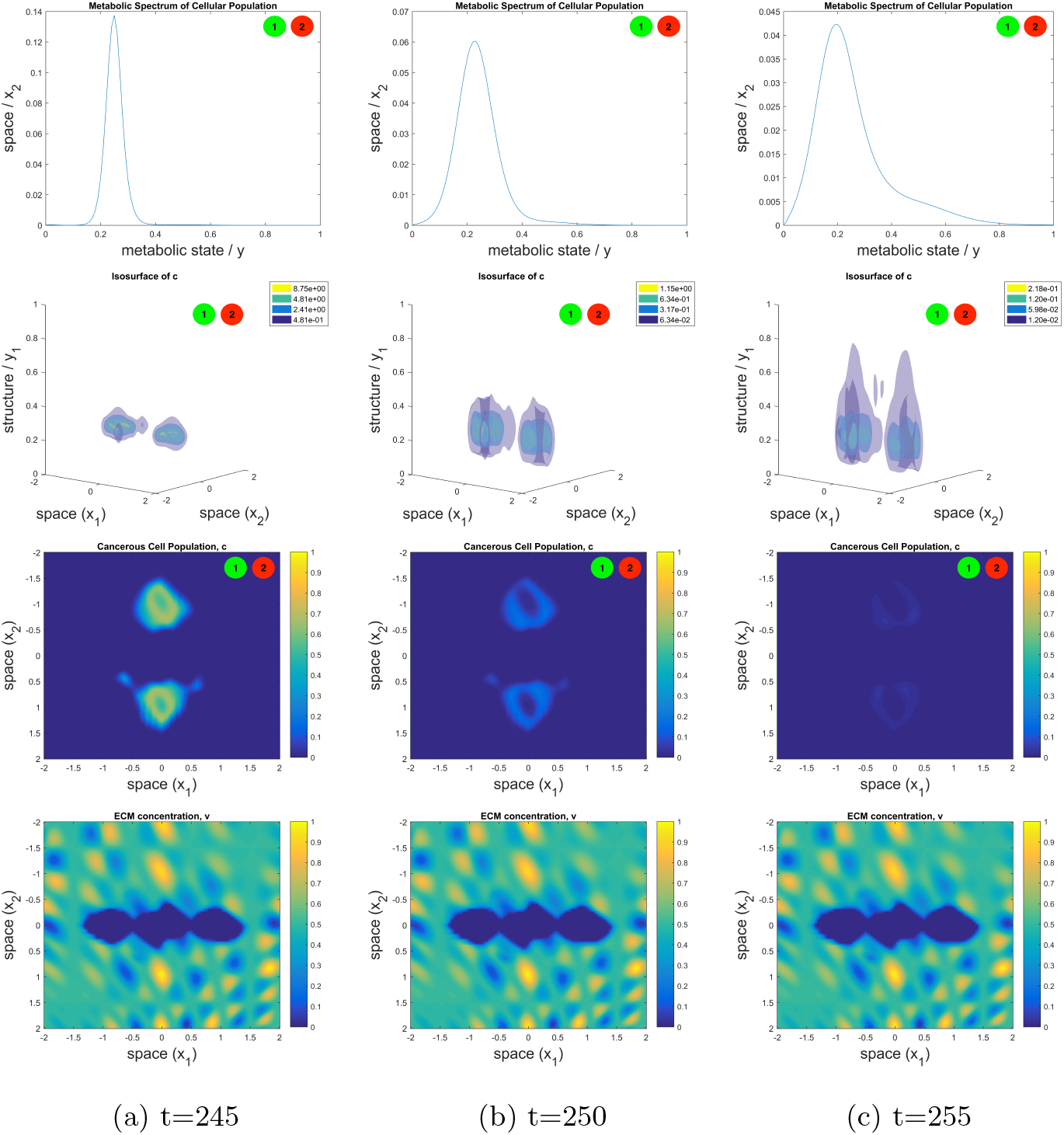
Tumours use oxphos metabolic pathways to resist targeted inhibition of glycolytic pathways by BRAFi and MEKi therapies. Shown are the phenotypic distribution (*1^st^ row*); the spatio-phenotypic surface distributions (*2^nd^ row*); and spatial distribution (*3^rd^ row*) of the cellular population. The spatial distribution of the ECNE, with colour-bar, is also shown (*4^th^ row*), for completeness and in order that one can place the tumour within its environmental context. All figures are given at times *t ∈* {245, 250, 255} within subfigures (a), (b), and (c) respectively. Within the surface plots, the colours represent surfaces of approximately equal concentrations within the spatio-phenotypic context of the cell gradiated from lowest to highest concentration as purple, blue, green, then yellow.

**Figure 15:**
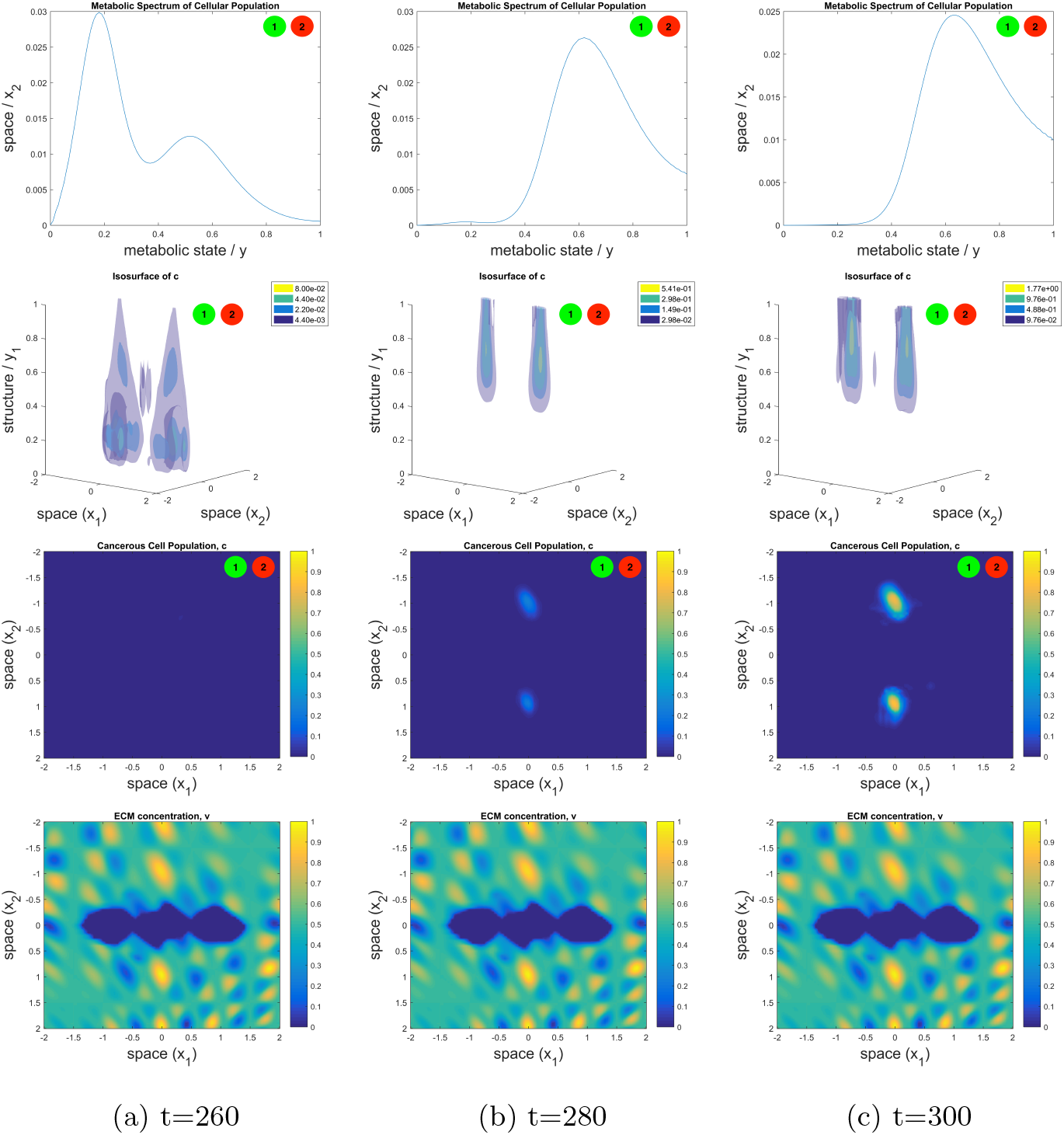
Tumours use oxphos metabolic pathways to resist targeted inhibition of glycolytic pathways by BRAFi and MEKi therapies. Shown are the phenotypic distribution (*1^st^ row*); the spatio-phenotypic surface distributions (*2^nd^ row*); and spatial distribution (*3^rd^ row*) of the cellular population. The spatial distribution of the ECNE, with colour-bar, is also shown (*4^th^ row*), for completeness and in order that one can place the tumour within its environmental context. All figures are given at times *t ∈* {260, 280, 300} within subfigures (a), (b), and (c) respectively. Within the surface plots, the colours represent surfaces of approximately equal concentrations within the spatio-phenotypic context of the cell gradiated from lowest to highest concentration as purple, blue, green, then yellow.

Observe, then, that in the initial growth phase (Fig. 12a) the cell population is tightly associated with a glycolytic metabolic state and that its spatial composition is compact, whilst during the sensitivity phase (Fig. 12b) the cell population begins to diverge from this behaviour and cells may be spatially observed further afield. Moreover, and throughout this phase, one can observe the degeneration of the narrow peak, during the initial growth phase (Fig. 12a, 1*^st^ row*), into a larger metabolic distribution centred at the same position as this initial peak (Fig. 12b, 1*^st^ row*). The increase in variance of the metabolic distribution is as a result of the diversification of metabolism under stressed conditions, whereas the displacement of the mean towards a resistant oxphos population (Fig. 12c, 1*^st^ row*) is as a result of selective pressure.

During the resistance phase, the newly oxphos population continues to proliferate (Fig. 13a), whilst any glycolytic cells are induced to apoptosis. When the drugs are washed from the tumour, however, at *t* = 210 one observes the cellular population beginning to migrate monotonically towards its preferred metabolic state (Fig. 13b, 1*^st^ & 2^nd^ rows*), *ω_c_* as observed at earlier time points (Fig. 12a, 1*^st^ row*), before reestablishing its glycolytic phenotype *y* ≈ *ω_c_* = 0.2 at *t* = 240 (Fig. 13c). This whole process is then repeated during the second wave of treatment (Fig. 13c, 14 & 15), with the tumour being visibly eradicated during a process of metabolic diversification and upheaval (Fig. 14b, 14c & 15a) before regrowing as an oxphos oriented tumour (Fig. 15b & 15c).

In this model, one may far more clearly see that the regrowth in the tumour is spatially correlated with the regions of highest ECNE concentrations (Fig. 12c, 3*^rd^ &* 4*^th^ rows*) and those regions where the cellular species will necessarily have the greatest access to nutrients. Interestingly, this will also be the spatial subregion in which the selective pressure is most elevated due to the presence of high concentrations of BRAFi+MEKi leading to the apoptosis of glycolytic cells and selecting for a more oxphos-dependent population of cells (Fig. 12c & 13a, 2*^nd^ row*).

To sum the above analysis of these results, the tumour exhibits an initially glycolytic mode of metabolism which, through stress-induced diversification, decays into a less defined mode of glucose metabolism. By spatially correlating with regions of heightened nutritional content, these resistant oxphos cells are able to outgrow their drug-induced apoptotic rate and proliferate. By removing the drug from the tumour, and the stressor of the cell, the cellular population attempts to reconsolidate its glycolytic state and increases its proliferative rate, ultimately allowing the second wave of treatment to visibly eradicate the remaining population of cells. Nevertheless, these cells are able to regain their metabolic advantage and return to an oxphos state, in order to once again become resistant to treatment.

### 6.3. More rapid secondary resistance wave may be explained by residual oxphos populations

One feature of the growth, which is of great clinical significance, is that of the increased rapidity to resistance upon the second wave of treatment (Fig. 11). In order to understand this, notice the pattern of metabolic migration in the cancer cell population, towards the preferred glycolytic state, during the drug holiday (Fig. 13). The tail on the right-hand side of the oxphos cell distribution (Fig. 13a & 13b, 1*^st^ row*) are not entirely consolidated during their backwards migration but, rather, remain as a residual oxphos cell population (Fig. 13b, 1*^st^ row*), which begin to appear upon selective degradation of glycolytic populations (Fig. 14b, 1*^st^ row*). Although these cells will migrate gradually towards their preferred metabolic state, *ω_c_*, it could be that their lower local nutritional value is allowing them to retain their oxphos state to a greater extent than the remainder of the population. Under the selective pressure applied by the drug, the glycolytic subpopulation is degraded, as it again attempts to diversify its metabolic status, whilst the oxphos population is free to grow (Fig. 14b, 14c, & 15, 1*st &* 2*^nd^ rows*), eventually replacing the glycolytic population as the dominant population within the tumour (Fig. 15b & 15c).

One may also clearly observe the difference in the spatio-structural distributions 20 days posterior to the first wave of treatment (Fig. 12b, 2*^nd^ row*) in comparison to 20 days posterior to the second (Fig. 15a, 2*^nd^ row*). After the first wave of treatment, the tumour having never been exposed to stress prior to this event, the metabolic profile of the tumour is neatly distributed around its preferred glycolytic state. After the second wave of treatment, however, the metabolic profile is bimodal, with a distinct oxphos as well as a glycolytic population. This appears to be due to the fact that not all of the cells from the resistant oxphos population have migrated fully back to their preferred glycolytic state and are, thus, able to repopulate the new resistant population far more rapidly since they are not subject to the same selective pressures as their glycolytic counterparts.

## 7. Discussion

We have introduced a general modelling framework for evolution of heterogeneity in solid tumours submitted to multiple drug therapy, wherein the definition of an appropriate normalized structural velocity, Ψ(*y*, *m̄*, *p̄*); structural diffusion matrix, Σ(*y*, *m̄*, *p̄*); growth function, Φ_*c*_(*y*, *m̄*, *c*, *υ*); and vector valued drug effectiveness function, *f̄* (*y*), may give rise to importantly nuanced patterns of behaviour. Using this framework, we then introduced two primary models for considering different dynamics within a tumour population. Firstly, the mutational model considered population level dynamics for a system in which an individual cell will sequentially undergo a BRAF mutation, followed by subsequent mutations which confer resistance to BRAFi and ipilimumab therapies. Secondly, we considered a plastic model of drug resistance, in which the switching of cellular dependence on glycolytic and oxphos pathways for the metabolism of glucose may confer a survival advantage when faced with glycolysis inhibiting BRAFi+MEKi treatments.

Using our mutational model to consider paradigms of punctuated equilibrium and phyletic gradualism in the evolution of the cellular genome, we found that punctuated equilibrium assumptions were more consistent with biological data. This shows good consistency with the modern cancer genomic literature, in asserting that short term catastrophes, rather than the gradual accumulation of mutations, is more likely to contribute to the mutational state of tumours [47, 48, 49]. We also predicted that using ipilimumab, immune cell-enhancers, in advance of a BRAFi is more effective at reducing the tumour population over the long term. This model prediction is confirmed by studies which used both ipilimumab and BRAFi [26].

Performing experiments for which drug was applied periodically in time we were able to qualitatively recapitulate the results of Thakur *et al.* [27]. We have suggested a mechanism for the apparently counterintuitive result of this experiment, that consists in keeping the tumor under control without completely eliminating the resistant subpopulation. We suggest that relative success of this therapy protocol in some tumours may imply their lesser mutated states at the initiation of the experiment, where the irregularity of the oscillations appears to depend on the number of different clones within or the clonal heterogeneity of the sample. This hypothesis may, presumably, be tested biologically in order to confirm this prediction from our model. The decay of the success of varying treatment strategies within a heterogeneous ECNE is consistent with the *in vivo* failure of treatments to adequately deal with tumours on the long term, and our experiments still predicted the preservation of the characteristic death and growth curves [22] under heterogeneous initial conditions.

Turning to the plastic metabolic model for the development of resistance to targeted therapies, we proposed and conformed the ability of such a model to predict re-sensitisation *in silico*. This model may then provide a clinical opportunity to model the success of therapy against such tumours on the basis of their respective environments (i.e. for tumours in differing tissue elasticities or densities). Moreover, our model illustrates the metabolic switching of the tumour as a continually heterogeneous spatio-structural population, allowing one to understand how spatial effects may influence structural resistance manoeuvres. The evolution of the glycolytic tumour to a metabolically oxphos cell populations, in combination with the coincidence of strongly selected cell populations and nutrient populations, may allow for the resistant proliferation of these subpopulations. These metabolically resistant populations will then preferentially re-sensitise themselves through metabolic remodeling, allowing for the effective second wave of treatment.

Moreover, our model provides an opportunity to understand the underlying dynamics of such metabolically plastic tumours and also the mechanisms of resistance and re-sensitisation, showing strong agreement with *in vivo* PDX tumour experiments. For both waves of treatment, our model shows a characteristic death, tolerance, and regrowth pattern, but with a quicker relapse occurring with the second wave of treatment. Experiments conducted by Ram-bow *et al.* [66] also show this pattern of death and growth, with faster regrowth posterior to the second wave of treatment, such that our model may provide an explanation of this phenomenon. Residual, metabolically resistant cells from the first wave of treatment may provide a basis for a resistant population to grow back more quickly upon the second wave of treatment. Implicitly, our model would predict that reducing treatment to as great an extent as is possible, whilst still eradicating the tumour, would reduce the opportunity for the tumour to establish this residual population and resist future waves of treatment.

## Appendix A. Thorough Discussion and Justification of the Mathematical Model

Current modelling approaches consider the cell as a discretely changing variable who exists in an explicitly sensitive or resistant state. We wish, here, to conceive of the cell as a continuously changing and finely tunable evolutionary population. Different cancer cells have similar, if not identical, origins and are not innately differentiable but have rather gained different characteristics. Therefore, we introduce a novel modelling framework in order to reconceive the mathematical representation of the cell, from this more nuanced perspective.

Cells do, however, function differently. Within these categories, then, there must be a wealth of diversity to reflect the reality of the structural differences between cells. In order to reflect this, we incorporate a term that operates similarly to those structural models previously employed [5, 6], whilst building on the solid mathematical derivation given by existing spatio-structuro-temporal models [9, 11]. Letting *ℐ*:= [0, *T*] ⊂ ℝ_+_ be the time interval over which the experiment is conducted; 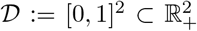 be the spatial domain; and 𝒫:= [0, 1] ⊂ ℝ_+_ define the continuous domain over which the mutational or metabolic changes may occur, we couple these dynamics using a simple conservation of mass assumption. If *V × W ⊆ 𝒟 × 𝒫* is an arbitrary volume of the spatio-structural domain with piecewise smooth boundaries *∂V* and *∂W* respectively, then we can write that the total population of cells in this volume is given by

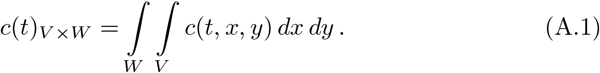

Then we can use an existing mathematical framework [9, 11] to deduce that the change in cell density *c*(*t*, *x*, *y*) is given by the partial differential equation

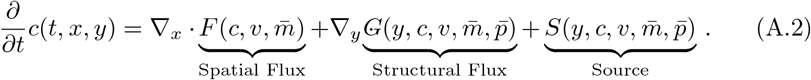

Let Ψ(*y*, *m̄*, *p̄*): *ℐ×𝓓 ×𝒫 →* ℝ be the normalized structural velocity for the cellular population. During a time interval of small length Δ*t*, those cells having the mutational or metabolic state *y* initially at *t*, will evolve to a state *y* + *r_μ_*Ψ(*y*, *m̄*, *p̄*)Δ*t* at *t* + Δ*t*, where *r_μ_* is the mean mutation rate. Moreover, let Σ(*y*, *m̄*, *p̄*): *ℐ × 𝓓 ×𝒫* be the structural diffusion matrix for the cellular population. Hence, the structural flux reads

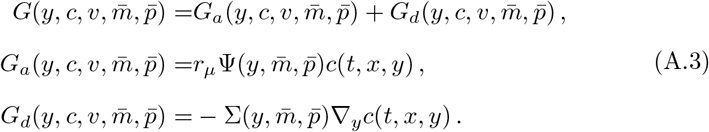

With this concept of a continuum of phenotypic progression, we then recognise that pharmaceuticals are generally targeted at specific metabolic pathways (related to selected cancer-related phenotypes and their respectively triggered mechanisms). Therefore, we employ a description of a phenotypic ‘spectrum’ wherein cells may inhabit any point on that available spectrum in *y*. These drugs may then target specific regions on this spectrum which employ the molecular pathways inhibited by these drugs. For this we form an effectiveness vector

*f̄* (*y*) ∈ *𝑦^P^* which describes the bandwidth in the mutational dimension *P* on which the drug is effective at diminishing the population of cells, for each given drug, *p_j_, j* ∈ {0, …, *P* }.

### Appendix A.1. Discussion on Single-Dosage Systems

The choice of mutational rate of change function would have to accurately represent the most sensible possible case for PG and PE, respectively. From (A.3) it is clear that the no-flux boundary condition is fulfilled automatically if the structural veolocity satisfies

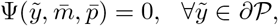

where *∂𝒫* is the boundary of the structural domain *𝒫*.

The structural velocity for PG is considered to be constant, except for a small region at the boundary. In order to construct such a function, we start with

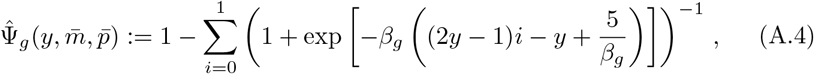

where *β_g_* is chosen sufficiently large such that the function Ψ̂*_g_* (*y*, *m̄*, *p̄*) is close to one everywhere except at narrow neighborhoods of *y* = 0 and of *y* = 1. The symmetry of the function Ψ̂_*g*_ implies that no-flux boundary conditions can be achieved by the imposition of

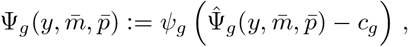

where the lower case *ψ_g_* gives the mutation rate parameter and *c_g_*:= Ψ̂_*g*_ (0, *m̄*, *p̄*) = Ψ̂_*g*_ (1, *m̄*, *p̄*) (Fig. 2a).

For the PE function, one must consider several features. Beyond smoothness, that is needed for both technical and biological reasons, one must again satisfy the no-flux conditions and impose the further conditions

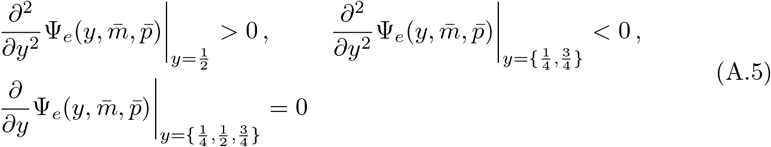

which is to say that maximal mutational velocity should occur between points of phyletic stability, “equilibria”, and minimal velocity should occur at intermediate points of phyletic stability (where boundary conditions cover the cases of minimal and maximal phyletic deviance). Thus, one can choose a function of the form

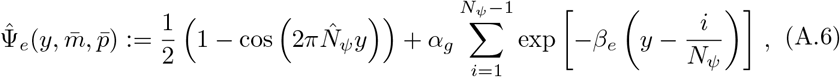

where *N_ψ_* = 3 is the number of absolute mutational states in the considered paradigm (pre-mutated, BRAF mutated, & resistantly mutated); *β_e_* is chosen such that distribution is increased smoothly; and the symmetry of this function in the domain implies that the no-flux boundary conditions can be satisfied by imposing

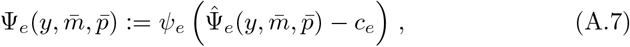

where *ψ_e_* again gives the mutational rate and *c_e_*:= Ψ̂_*e*_(0, *m̄*, *p̄*) = Ψ̂_*e*_(1, *m̄*, *p̄*) (Fig. 2b).

Defining the cancerous population as being represented by a continuous distribution in the mutational space further allows one to define the drug effectiveness functions such that the drugs themselves target, not a discrete subset of the cellular population but rather, a continuous distribution of phenotypes which correspond to states in *𝒫*. With this in mind, we begin by defining the function itself as being represented by a vector of such distributions, each distribution describing the action of an individual drug

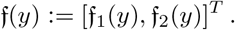

Then, we can continue by defining the individual effectiveness functions of each drug as being given by the standard Gaussian distribution in *y*, such that

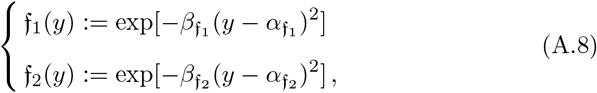

where 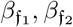 define, reciprocally, the breadth of the distribution and 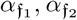 define the mutational position, in *y*, of maximal effectiveness.

In this case, since we consider *N_ψ_* = 3 mutationally-equidistant resting states for the tumour population, we define that the BRAF inhibitor, *p*_1_, targets a mutational state which corresponds to the most susceptible of these resting (low relative mutational rate) states, 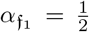. In this sense, also, we assert that the resting state, 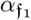, represents a mutational state wherein the cell has a BRAF mutation, which makes cells in this state most susceptible to BRAFi treatment.

We then make a phenomenological choice, although with some support from the existing literature, to maximise the effectiveness of drug *p*_2_, i.e. ipilimumab, at an arbitrary position between the establishment of mutational states corresponding to 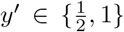, such that 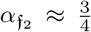. We interpret the choice of 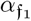 (smaller than 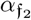) as conveying the sense that BRAFi is maximally responsive at early mutational states where PTEN mutation is developing but not established within the population, whereas ipilumab is maximally responsive at later stages. We assume this on the basis that treatment with BRAFi, prior to treatment with immune cell enhancers, is ineffective as opposed to the contrary and that this implies that sensitivity to BRAFi may occur at an early stage of mutational development.

### Appendix A.2. Discussion on Multi-Dosage Systems

Thusly, we describe the metabolic change function, Ψ: *𝒟×𝒫* → ℝ, in terms of the phenotypic stress on the cell. We assume, firstly, that under a condition in which the influence of stressors is minimised, the cell has a preferred phenotypic state at *y* = *ω_c_*, which corresponds to a given utilisation of each pathway. We also assume that the primary stressors for the cell are malnutrition, which will be a function of *m*_2_, the presence of BRAFi, *p*_1_, and that of MEKi, *p*_2_, which act to deplete the cells ability to proliferate effectively.

Then the non-stressed term in the function must be given such that phenotypic advection is positive below this preferential state and negative above this state such that it will depend upon the relation 1 − *ω_c_* for a non-dimensionalised system. The non-stressed condition must then be given by the opposing probability to that of stress such that 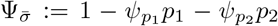, with parameters chosen such that Ψ_*σ̄*_ ≥ 0, *∀*(*t*, *x*) ∈ *ℐ×𝒟*.

Stressed conditions for the cell are then quantified by the gradient of the cellular concentration in the region, giving a measurement of the collectivity of the behaviours of local cells. This choice of function for stressed conditions gives rise to diffusion under cellular stress, the rationale for which can be given by the intuitive understanding that cells diversify their behaviours in the presence of stressors. The magnitude of this stress is then determined by the concentrations of BRAFi, *p*_1_, and MEKi, *p*_2_, and is linearly diminished with the concentration of nutrient species, *m*_2_. All of these factors act as stressors to the cell and have their relative effects quantified by the weights 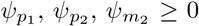, respectively. Then, the structural flux has diffusion and advection terms as follows

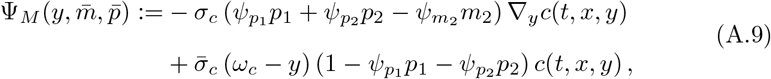

with the introduction of the stress, *σ_c_*, and non-stress, *σ̄_c_*, parameters determining the weightings of the diffusion and advection terms with respect to one another.

Now, one must consider the nature and form of the effectiveness functions for the drug species, BRAFi (*p*_1_) and MEKi (*p*_2_), on the cellular population, in terms of their effect on the glycolytic or oxphos pathways. Firstly, we begin by writing the vector

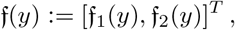

to represent functions *f*_1_(*y*) and *f*_2_(*y*) in compact notation and begin by noticing that both of these drugs target genes essential to glycolysis. The transcription factors HIF1*α*, c-Myc, and Mondo A have been found to be downstream upregulators of glycolytic behaviours in BRAF^υ600^ cells [67, 71]. Moreover, BRAFi has been shown to prevent the hyperswitching of mutant melanoma cells to pyruvate based metabolism [72] – the primary product of glycolysis.

Withal, MEKi is responsible for targeting this same pathway, in melanoma cells. It has also been found that the PI3K pathway, activated by MEK, is responsible for glucose transport, and glycolytic metabolism, and can be inhibited by inhibition of MEK [73, 74].

The biological literature points to a link between melanoma associated genes, including BRAF and MEK, and the glycolytic pathway for glucose metabolism. Therefore, we write that the standard forms of the effectiveness functions will be Gaussian functions, with low values for variance, or high values for 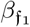 and 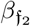, such that

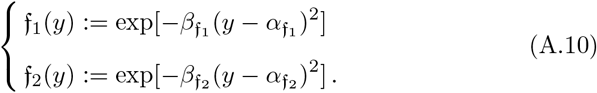

The values around which these functions are centred, 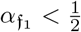 and 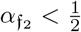, are chosen to align with the peak effect of the drug on the glycolytic and oxphos pathways.

Finally, we choose the proliferation function, *ϕ_M_*: *ℐ×𝒟 ×𝒫 →* ℝ, such that it is space-wise logistic in *c*(*t*, *x*, *y*). Moreover, we assume that the cellular population requires nutrient in order to achieve positive proliferation and choose some arbitrary threshold value 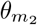 in order that, below such a value, the cellular population is depleted due to malnutrition. It is then imposed upon the system that there are two concurrent modes of proliferation: glycolytic and non-glycolytic. The non-glycolytic mode is not dependent upon the phenotypic state of the cell, *y*, and is rather an underlying process of all cells, whereas the glycolytic pathway is linearly enhanced by the percentage of glycolytic metabolism utilised (such that it is maximal at *y* = 0). This is justified on account of the excess lipids produced through utilisation of glycolytic pathways. Therefore, we write

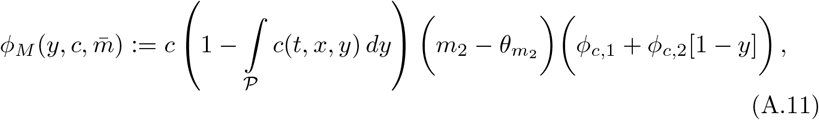

where 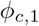 and 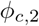 give the rates of non-glycolytic and glycolytic metabolism, respectively.

**Table B.1:** List of parameters used for numerical simulations of the model. Parameters are defined within a non-dimensionalised system (excepting for time measured in days) and, as such, are defined in terms of units days^−1^.

| | $c$ | $v$ | $m_1$ | $m_2$ | $p$ |
| --- | --- | --- | --- | --- | --- |
| $D$ | $1 \times 10^{-5}$ | | $1 \times 10^{-4}$ | $1 \times 10^{-4}$ | $5 \times 10^{-4}$ |
| $\phi$ | 0.4 | $5 \times 10^{-2}$ | 0.1 | 0.1 | |
| $\delta$ | 2 | 10 | 0.1 | $5 \times 10^{-2}$ | $5 \times 10^{-2}$ |
| $\chi$ | | $5 \times 10^{-5}$ | $1 \times 10^{-3}$ | 0 | 0 |

## Appendix B. Numerical Methods

### Appendix B.1. Methods for Mutational, Single-Dosage System

Initial conditions were chosen to be consistent with previous models [53] and for consistency with the biological methodology, as regards the impregnation of mice with cancerous cells. The particular study, using animal models, with which we compare our results injected mice with approximately 5 × 10^3^ − 2 × 10^5^ cells [75]. Therefore, our initial conditions reflect this with

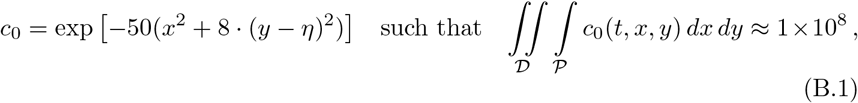

where, since we know that the biological experiments were initiated with an approximate cell count of 2.5 × 10^4^ cells, we assume that the cellular distribution is measured approximately in 10^3^ 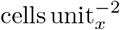. Further, the default initial location in the phenotypic dimension is given by 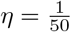. One should also clarify that this constitutes not an entirely pre-mutated cell population but an already heterogeneous mixture of cells with at least one precursor event that induces the early stages of the BRAF mutation process.

Other quantities for which it is imperative that one have measures include the gross spatial population, which is given by the cellular population taken over the entirety of the structure domain, *𝒫*, and is given by

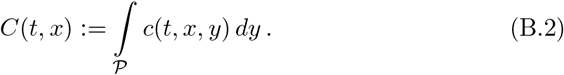

To calculate the volume of their tumour from its 2-dimensional section, Perna *et al*. [22] measure the lengths of the major and minor axes of the visible tumour, given by *a* and *b* respectively, and use the formula of an ellipsoid to write

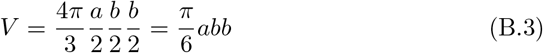

In order to avoid having to define the value of our function, *c*(*t, x, y*), above which the tumour would constitute a visible tumour, which would otherwise be given by a threshold of visibility *θ_υ_*, we assume the proportionality of the tumour mass and the area of the section over which the tumour is visible, written as

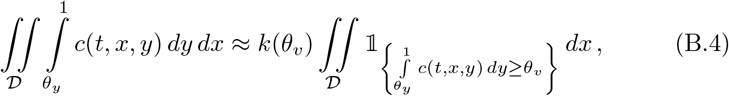

where the proportionality constant is dependent on the visibility threshold and is given by *k*: ℝ *→* ℝ. To calculate the model’s tumour volume, i.e. the volume of cells which have developed into cancerous subtypes, we then take the mass of the tumour at *y ≥ θ_y_* and invoke the calculation from the tuning model [22] such that

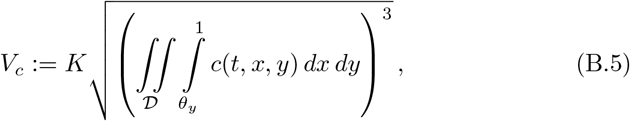

with the adaptation of the ellipsoidal volume equation to 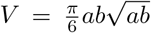 and where we take that *θ_y_* = 0.2 and *K* is an arbitrary constant.

Then, in order to carry out our test experiment, we control the heterogeneity using the following formula for the initial condition
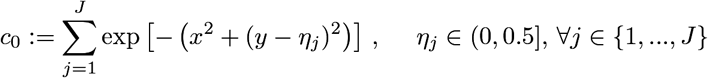

and also in line with the initial volume condition (B.1), and where *J* is in some sense a measure of the initial heterogeneity. We then apply the drug dosage periodically in time intervals given by [0, 20] ∪ [40, 60] ∪ [80, 100] ∪ [120, 140]. For the simulations given in this current study, we use the range *J* ∈ {1*, …,* 5} to establish example data.

### Appendix B.2. Methods for Metabolic, Multi-Dosage Systems

Due to the nature of the structural flux (A.9), it is necessary to develop a set of zero-flux boundary conditions which prevent, for example, diffusion in *y* from causing cells to exit the domain, *𝒫*. Although (A.9) has both advection and diffusion terms, the metabolic change function is defined such that Ψ_*σ̄*_(*y*) = 0*, y* ∈ *∂𝒫*, meaning that advection fluxes are identically zero on the boundary. Therefore, we simply implement zero-Neumann boundary conditions on structural diffusion fluxes, namely ∇*_y_c*(*t, x, y*) = 0*, y* ∈ *∂𝒫*.

To begin treatment, one gradual dosage was given between *t* = 80 and *t* = 100, linearly in time, *t*. The drug was then washed from the tumour, in a step-wise fashion, at *t* = 210, as this is the point at which the tumour volume had regrown to ∼20% of its previous maximum, and the tumour was allowed to regrow, unencumbered by glycolytic inhibitors for 30 days. A second gradual dosage was then given between *t* = 240 and *t* = 260, whereafter no further interventions were made.

Further, we define the unique structured population profile by the cellular population over the entirety of the spatial domain, *D*, given by

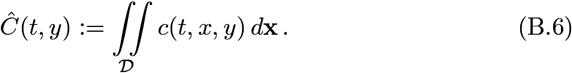

This can be used to describe the metabolic or structural profile of the tumour at a given time, *t*.

## Acknowledgements

The authors gratefully acknowledge financial support from the Canceropole GSO and École Doctorale I2S de l’Université de Montpellier. We are also grateful to N. Theret, A. Devenyi, and D. Fisher for their critical reading of the manuscript.

